# Enhanced inflammatory signaling driven by metabolic switch in Aicardi-Goutières syndrome

**DOI:** 10.1101/2023.02.23.529707

**Authors:** Maxime Batignes, Marine Luka, Surabhi Jagtap, Camille de Cevins, Ivan Nemazanyy, Tinhinane Fali, Víctor García-Paredes, Francesco Carbone, Brieuc P. Pérot, Bénédicte Neven, Brigitte Bader-Meunier, Pierre Quartier dit Maire, Marie Hully, Alexandre Belot, Alice Lepelley, Marie-Louise Frémond, Yanick J. Crow, Alain Fischer, Mickaël M. Ménager

## Abstract

Aicardi-Goutières syndrome (AGS) is a genetic type I interferon (IFN)-mediated disease characterised by neurological involvement with onset in childhood. Chronic inflammation in response to uncontrolled type I IFN production is, among other things, associated with IP-10 secretion. We analysed, at the single-cell transcriptomic levels, peripheral blood samples from patients bearing AGS-causing mutations in *SAMHD1*, *RNASEH2B* or *ADAR1* genes. Using machine-learning approaches and differential gene expression we identified a drastic loss of transcription factor hypoxia induced factor 1 α (HIF-1α) expression and activity associated with features of a metabolic switch and mitochondrial stress in monocytes/dendritic cells. Chemical stabilization of HIF-1α, with a synthetic drug in an *in vitro* model of AGS, allowed us to reverse the energy metabolic switch, attenuate mitochondrial stress and markedly reduce IP-10 production. We therefore propose that energy metabolic switch contributes to exacerbated chronic inflammation in AGS, and that targeting this pathway might represent a promising therapeutic approach.

## Introduction

Aicardi-Goutières syndrome (AGS) is a monogenic disorder with onset frequently observed in the first months of life. Mutations causing AGS have been identified in genes encoding for the exonuclease TREX1 (AGS1) ^1^, the components B, C and A of the RNase H2 complex (RNASEH2B (AGS2), RNASEH2C (AGS3), RNASEH2A (AGS4)) ^2^, the deoxynucleotide triphosphate triphosphohydrolase SAMHD1 (AGS5) ^3^, the double-stranded RNA-specific adenosine deaminase (ADAR1 (AGS6)) ^4^, the cytosolic dsRNA sensor IFIH1 (AGS7) ^5^ and components of the replication-dependent histone pre-mRNA-processing complex LSM11 (AGS8) and RNU7-1 (AGS9) ^6^. The encoded proteins are involved in the processing (AGS1- 6) or sensing (AGS7) of cellular nucleic acid, and disruption of their normal function can induce an aberrant antiviral transcriptomic response ^7^. These pathways can be separated into two arms: on one hand, detection of dsDNA and RNA/DNA hybrid leading to the activation of the stimulator of interferon genes (STING) pathway; and on the other hand, dsRNA sensing by the mitochondrial antiviral-signaling protein (MAVS). Both STING and MAVS activation converge on the activation by phosphorylation and nuclear translocation of the transcription factor IRF3, leading to the production of type I IFN and the strengthening of cell defence mechanisms against viruses ^8^. Mutations described in AGS lead to constitutive triggering of the nucleic acid sensing pathways and the production of type I IFN, even in the absence of viral infection ^6, 9^. Thus, AGS is a chronic, inflammatory disorder that can mimics *in utero*-acquired viral infection but differs, notably, in the type and nature of cytokines produced. In AGS, the inflammatory response is orchestrated by type I IFN signaling but also involves inflammatory signals through IFN-induced protein 10 (IP-10, also called CXCL10), CCL2 and vascular endothelial growth factor (VEGF), with no obvious upregulation of IL-6 or IL-8 ^10^, typically observed in response to viral infections. The major clinical features of AGS relate to the brain, most particularly intracranial calcification, white matter disease and cerebral atrophy. Extra-neurological features can also occur, including vasculitic chilblain-like lesions, and features consistent with systemic lupus erythematosus (SLE) in some cases ^9^. While type I IFN levels can be difficult to measure in patients, its activity can be monitored by following the expression of IFN-stimulated genes (ISG) such as IP-10, which concentration levels are consistently elevated both in serum and the cerebrospinal fluid of patients ^10^. Importantly, the type I IFN response and IP-10 have been shown to induce direct neurotoxicity in humans ^11^. Of note, type I IFN targeted therapies have proven of limited impact on the main neurological features of AGS ^12^, arguing for a need to better understand cellular pathways involved in AGS. In the present study, we screened peripheral blood mononuclear cells (PBMC) from AGS patients using single cell RNA sequencing (scRNA-seq) to reveal potentially new cellular mechanisms relevant to the pathogenesis.

## Results

### Marked changes in the transcriptomic profile of monocytes/cDCs in AGS

PBMC from 6 unrelated patients with AGS and 4 unrelated healthy donors were extracted in two separated sets of experiments. Two experiments were performed on two patients harbouring biallelic mutations in *SAMHD1* gene (AGS5). scRNA-seq libraries for these two AGS5 patients (AGS5: P1 and P2) were generated and integrated with PBMC from three unrelated healthy individuals (CTRL: C1, C2 and C3) to generate integration#1 (**fig1A**). In a third experiment, we generated scRNA-seq libraries from two patients with biallelic mutations in *RNASEH2B* gene (AGS2: P3 and P4) and from two patients with mutations in *ADAR1* gene (AGS6: P5 and P6). We added data from two symptomatic patients bearing a dominant-negative mutation in *COPA* gene (COPA: P7 and P8). The COPA syndrome is mediated through chronic STING activation even in the absence of nucleic acid sensing ^13^. Thus, COPA syndrome is also a type I IFN-driven disease, although, in contrast to AGS patients, COPA manifests as chronic lung inflammation in the absence of overt neurological involvement. None of the patients in the second set of experiment were under immunosuppressive or reverse transcriptase inhibitors (RTI) therapy at the time of sampling. Finally, we used PBMC from 2 unrelated healthy individuals (CTRL: C4 and C5) to complete the generation of integration#2 (**fig1B**). As libraries generated in integrations #1 and #2 were performed with different chemical versions of 10X genomics technology, we analysed these samples separately to minimize any batch effect. A first uniform manifold approximation and projection (UMAP) was generated for integration#1 (**fig1C**), including patients harbouring *SAMHD1* mutations and healthy controls, and a second UMAP was generated for integration#2 (**fig1D**) with healthy controls and patients bearing mutations in *RNASEH2B*, *ADAR1* and *COPA* genes (**see supplementary information table SI1**).

**Fig 1:**
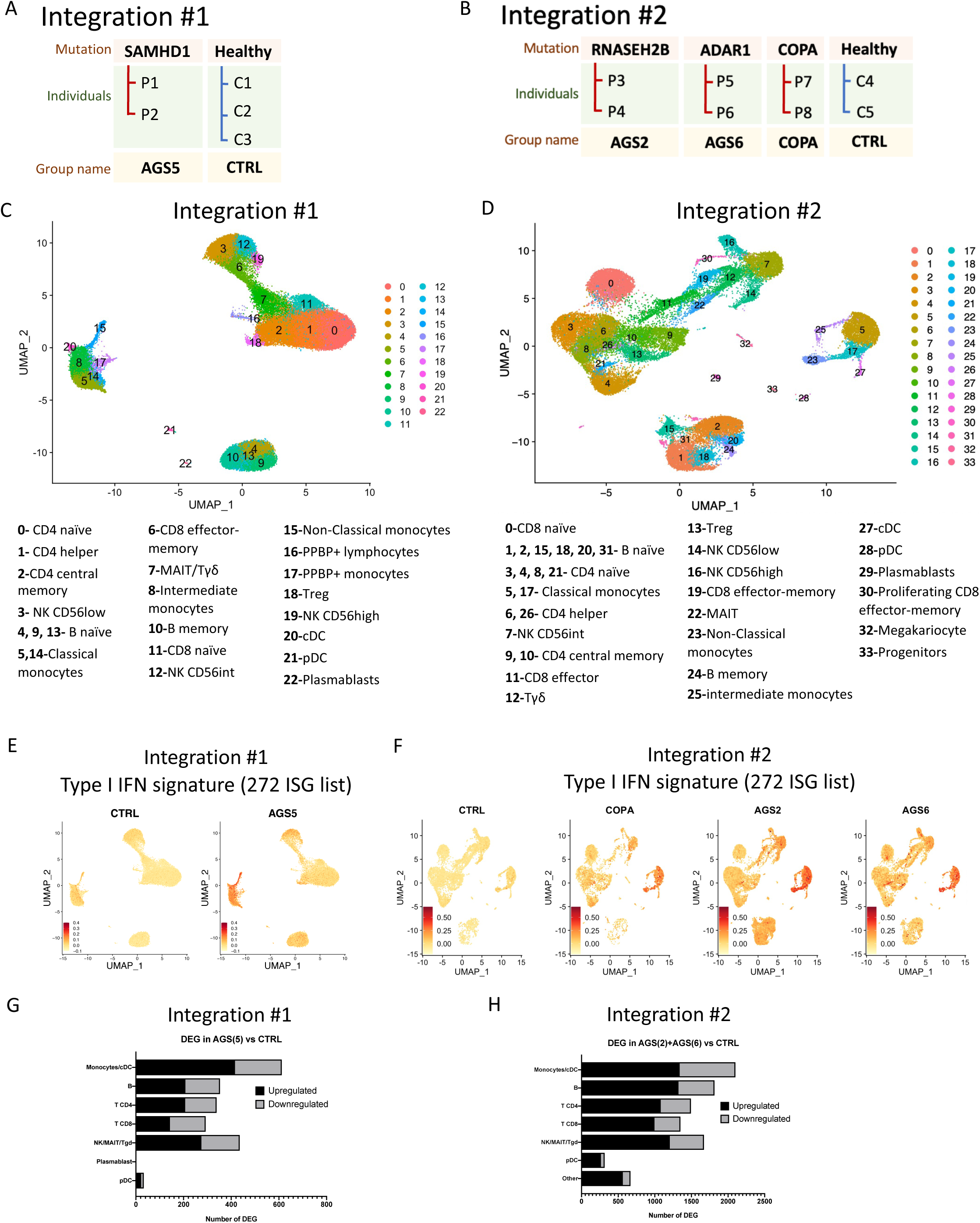
Drastic changes in transcriptomic profiles of monocytes/cDC during AGS. **(A and B)** Samples information and scRNA-seq integration strategy. Data from PBMC of two patients developing AGS with a mutation in *SAMHD1* gene (P1 and P2), were identified as group AGS5. Patients scRNA-seq data were integrated with data from PBMC of 3 unrelated healthy donors (CTRL group) to generate integration#1. P1 was under 5 month of reverse transcriptase inhibitor tri-therapy (RTI). P2 was sampled before and after 1 month of RTI treatment **(A)**. Data from PBMC of two patients with a mutation in *RNASEH2B* gene (P3 and P4), two patients with a mutation in *ADAR1* gene (P5 and P6) and two patients with a mutation in *COPA* gene (P7 and P8) were identified as AGS2, AGS6 and COPA groups, respectively. These unrelated samples were integrated with data from PBMC of two unrelated healthy controls (CTRL group) to generate integration#2. None of the samples in integration#2 were under treatment at time of sampling **(B)**. Integration#1 and #2 could not be merge due to differences in protocol used for library generation. **(C and D)** UMAP generated with Seurat at resolution 1.6 with cluster annotation of integration#1 **(C)** and #2 **(D)**. Gene markers used for cluster identification are listed in table SI2. **(E and F)** Features plot of ISG signature score of 272 ISG (from Rosenberg et al. 2018) in integration#1 **(E)** or integration#2 **(F)**. UMAP are split by CTRL, AGS2, AGS5, AGS6 and COPA groups. g=genes **(G and H)** Bar graph of the number of differentially expressed genes (DEG) in each cell types of integration#1 **(G)** and integration#2 **(H)** in AGS versus CTRL groups. In integration #1, cell types regroup several clusters at 1.6 resolution: “Monocytes/cDC” (clusters 5, 14, 17, 8, 20, 15), “B” (clusters: 4, 9, 10, 13,), “CD4” (clusters 0, 1, 2, 16, 18), “CD8” (clusters 6, 11), “NK/MAIT/Tγδ” (clusters 3, 7, 12, 19), pDC (cluster 21) and “Plasmablast” (cluster 22). In integration#2 cell types regroup several clusters at 1.6 resolution: “Monocytes/cDC” (clusters 5, 17, 23, 25, 27), “B” (clusters: 1, 2, 15, 18, 20, 24, 31), “CD4” (clusters 3, 4, 6, 8, 9, 10, 13, 21, 26), “CD8” (clusters 0, 11, 19, 30), “NK/MAIT/Tgd” (clusters 7, 12, 14, 16, 22), pDC (cluster 28) and “Others” (cluster 29, 32, 33). cDC= classical Dendritic cells; B= B lymphocytes; CD8= Lymphocytes T CD8^+^; CD4= Lymphocytes T CD4^+^; NK= Natural Killer cells; MAIT= Mucosal-associated invariant T cells; Tγδ= Lymphocytes Tγδ; pDC= plasmacytoid dendritic cells.

We first assessed cell type proportions in all samples. We detected a decreased proportion of effector memory T cells, NK cells, MAIT and γδ T cells in PBMC of AGS5 patients (**figS1A**). In parallel, the proportion of B cells was increased. We obtained comparable results for AGS2, AGS6 and COPA patients, except for NK CD56int, MAIT and CD8 effector memory cells, with proportions similar to control for AGS6 patients (**figS1B**).

As both AGS and COPA are type I IFN-related diseases, we then assessed IFN-stimulated gene (ISG) expression at the single-cell transcriptomic level. Specifically, we used a previously described signature of 272 ISG ^14^ and, in addition, we used a six-ISG “IFN signature” that has proven to be useful for diagnostic purposes in type I interferonopathies ^15^. We observed, as expected, an elevated type I IFN signature in AGS5 (**fig1E and S1C**), AGS2, AGS6 and COPA (**fig1F and S1D**) patients compared to respective controls. Interestingly, the strongest IFN response was observed in monocytes and classical dendritic cells (cDC). For most of the signature tested in this article, differences are less marked in integration#1 than in integration#2 due to a lower count of detected genes, but we systematically confirmed our observations by comparing the two integrations. Moreover, when differentially expressed genes (DEG) were extracted, it appeared that monocytes and cDC exhibited at least 40% more DEG between AGS patients and CTRL than other cells types in integration#1 and 15.7% more in integration#2 respectively (**fig1G and H**). This higher number of DEG in monocytes/cDC was not due to higher cell proportion in this group (**figS1F and G**). Thus, it suggests that greater transcriptomic changes occur in monocytes and cDC of AGS patients, although it should be mentioned that the highest number of genes detected by scRNA-seq was in monocytes and cDC (**figS1H and I**).

### Gene regulatory network analysis revealed a loss of HIF-1α transcription factor activity in PBMC of AGS patients

In order to gain an unsupervised understanding of factors driving the observed transcriptomic differences in AGS, we performed a gene regulatory network (GRN) analysis ^16^ applying pySCENIC (Single-Cell rEgulatory Network Inference and Clustering)^16^ on all cells from our scRNA-seq data sets. GRN allowed us to predict the transcription factors most likely to be important in the transcriptomic profile of CTRL PBMC and, then, of AGS patients. Briefly, co-expression modules containing genes and a putative TF were identified using GRNboost ^17^. TF binding motifs were evaluated in each gene of the module using RcisTarget. Only genes that present an enrichment for the binding motif of the module’s TF were kept in the final regulon. To explore those GRN, we decided to focus on the number of predicted outdegrees, which in our anlayses represents, the number of predicted targets inferred from the scRNA-seq data. Consequently, we were able to generate a network of TF and their predicted targets for CTRL group in integration#1 (**fig2A left**), AGS patients of integration #1 (**fig2A right**), CTRL group of integration#2 (**fig2C left**) and AGS patients of integration #2 (**fig2 C right**). We focused our analysis on the top 50 TF with the most predicted targets (termed out-degree) in the different GRN generated (**figS2A and S2B**). In order to estimate changes of activities of TF between CTRL and AGS patients, we then performed a differential out-degree analysis, by analyzing the ratio between the number of targets of each TF in AGS on the number of its targets in CTRL. Most of the TFs with increased out-degree between CTRL and AGS patients were related to inflammatory mechanisms and to type I IFN response such as *IRF6*, *IRF7* or *STAT1* (**fig2B and 2D**). In contrast, *HIF1A* stood out as the TF with the most important loss of target gene expressions in AGS compared to the CTRL group in integration#1 (**fig2B**) and integration#2 (**fig2D**), as also observed when looking at *HIF1A* in the different GRNs (fig 2A and 2C, red circles). Indeed, the number of predicted targets of *HIF1A* in AGS reached 0 in both integrations, while in CTRL of integration#1 and integration#2, it reached 82 and 163 respectively (**figS2A and S2B**). HIF-1α, encoded by *HIF1A*, is a sensor of energy metabolism stress and its transcriptional activity allows the adaptation of energy production pathways, orienting glucose consumption through anaerobic glycolysis instead of oxidative phosphorylation ^18, 19^. Overall, using an unsupervised approach with gene regulatory network analyses, we determined that in cell of AGS2, AGS5 and AGS6 patients, beside a previously described strong response to type I IFN and inflammation, a marked loss of HIF-1α activity was predicted, suggesting a potential role for this TF in AGS pathogenesis.

**Fig 2:**
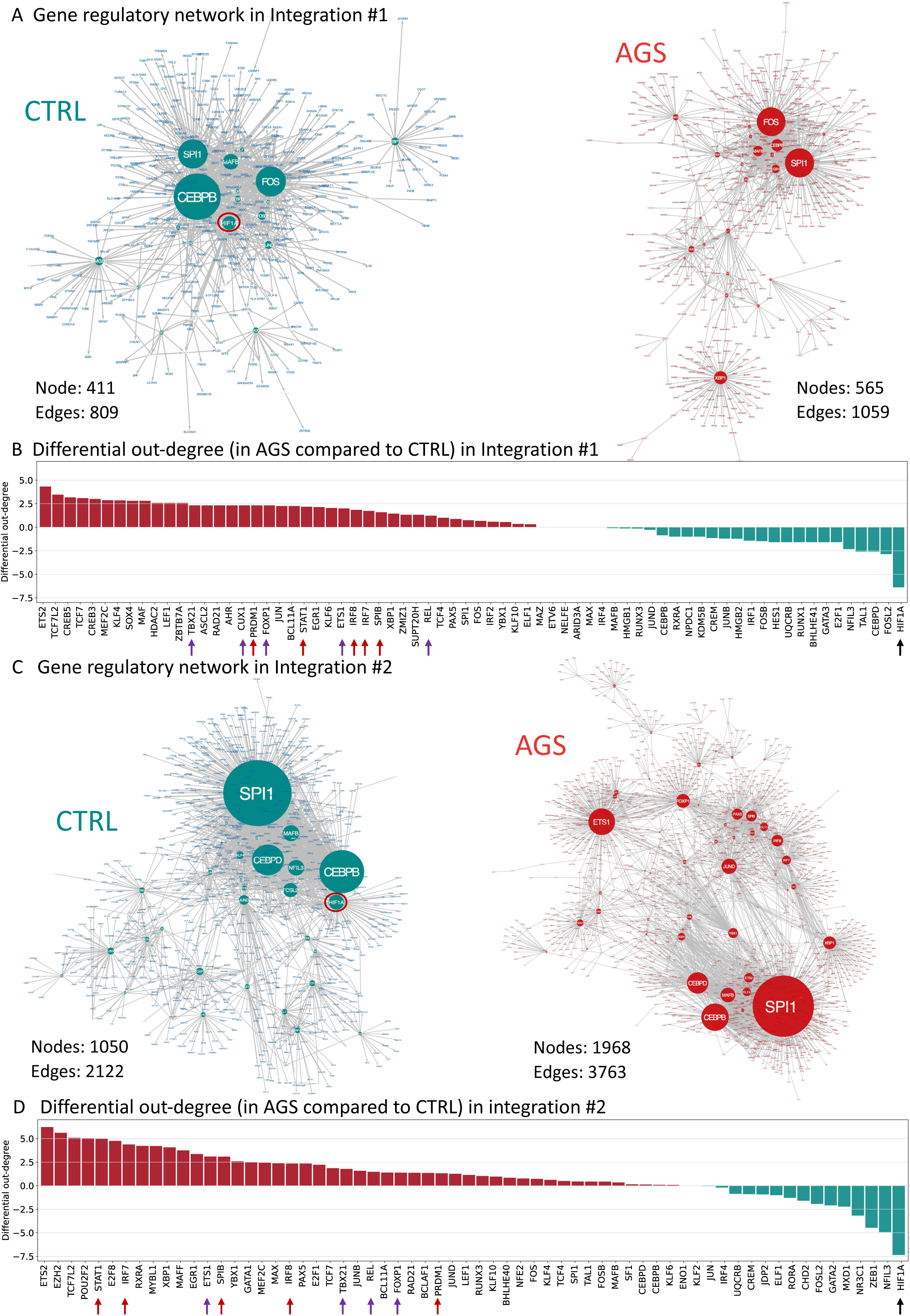
Gene regulatory network analyses predicted loss of HIF-1α activity in AGS patients. (A) Gene regulatory network on scRNA-seq data of integration#1 in PBMC from CTRL (C1, C2, C1bis, C3) (left panel) and from AGS patients (P1, P2_before_T, P2_before_T_bis, P2_after_T) (Right panel). Size of nodes indicates out-degree score (number of targets in scRNA-seq data predicted to be regulated by the transcription factor (TF)). Both TF and targets are displayed. Red circle highlights HIF1A TF. (B) Differential out degree analysis obtained by subtracting out-degree score of TF in AGS with corresponding out-degree score in CTRL for integration#1. Top50 TF with highest out- degree in CTRL are represented. Red bars highlight TF with increased out-degree score in AGS compared to CTRL while green bars highlight TF with decreased out-degree in patients. Red arrows point type I IFN related TF, purple arrows point inflammation related TF and black arrow points most important loss of out-degree in AGS. (C) Gene regulatory network on scRNA-seq data of integration#2 in PBMC from CTRL (C4 and C5) (left panel) and from all AGS patients (AGS2 + AGS6, i.e.: P3, P4, P5, P6) (Right panel). Size of nodes indicates the number of out degree (number of targets in scRNA-seq data predicted to be regulated by the TF). Both TF and targets are displayed. Red circle highlights HIF1A TF. (D) Differential out degree analysis obtained by subtracting out-degree score of TF in AGS with corresponding out-degree score in CTRL for integration#2. Top50 TF with highest out- degree in CTRL are represented. Red bars highlight TF with increased out-degree score in AGS compared to CTRL while green bars highlight TF with decreased out-degree in patients. Red arrows point type I IFN related TF, purple arrows point inflammation related TF and black arrow points most important loss of out-degree in AGS. See also figS2.

### Defective HIF-1α transcriptional program in monocytes/cDC of AGS patients

To validate and investigate the importance of the loss of HIF-1α-mediated transcriptional activity in AGS patients highlighted by our unsupervised approach, we decided to focus first on monocytes/cDC, where the highest response to type I IFN and number of dysregulated genes were detected (Fig 1). *HIF1A* appears among the most differentially downregulated genes in monocytes/cDC of both AGS5 (**fig3A**) and AGS2/AGS6 (**fig3B**) patients compared to CTRL. Indeed, *HIF1A* expression is decreased in monocytes/cDC of AGS patients (**fig3C and D**), and similarly in all subpopulations of monocytic-cell types (**figS3A and B**). Not only *HIF1A* expression level was lower, but the percentage of cells expressing detectable levels of *HIF1A* mRNA was also reduced in cells of AGS patient, particularly in monocytes/cDC compared to any other cell type (**figS3C, D, E and F**). Cells from COPA patient exhibited an intermediate level of *HIF1A* expression between CTRL and AGS2/AGS6 patients (**fig3C and D**). Yeh et al. first suggested that HIF-1α protein levels can be downregulated by ISG15-induced proteasome degradation in response to type I IFN ^20^. Thus, we plotted *HIF1A* expression against type I IFN signature in monocytes/cDC. Cells expressing high levels of type I IFN signature score expressed low levels of *HIF1A* (**fig3E and F**). However, the correlation was not strict, as monocytes/cDC with a low IFN signature demonstrated variable levels of *HIF1A* mRNA (**fig S3G and H**). These data suggest that enhanced type I IFN signaling in cell of AGS and COPA patients, might repress *HIF1A* expression at the mRNA level. To verify that lower *HIF1A* expression correlates with decreased HIF-1α transcriptomic activity, we generated a signature of known HIF-1α targets from the NCI-Natures database ^21^. Monocytes/cDC of AGS patients demonstrated a lower signature score of HIF-1α targets compared to CTRL (**fig3G and H**). Again, cells from COPA patients reached an intermediate level between CTRL and AGS2/AGS6 patients. Among all HIF-1α targets, vascular endothelial growth factor A (*VEGFA*) is one of the most sensitive targets to HIF-1α regulation ^22^. Following *HIF1A* expression, *VEGFA* was found mainly expressed in monocytes/cDC of CTRL, and completely lost in cells of patients (**fig3I and J**). We confirmed here a clear loss of *HIF1A* expression and transcriptional activity in AGS patients. Regarding COPA syndrome, patients present an intermediate *HIF1A* expression and transcriptional activity between CTRL individuals and AGS patients, corresponding to an intermediate type I IFN response.

**Fig 3:**
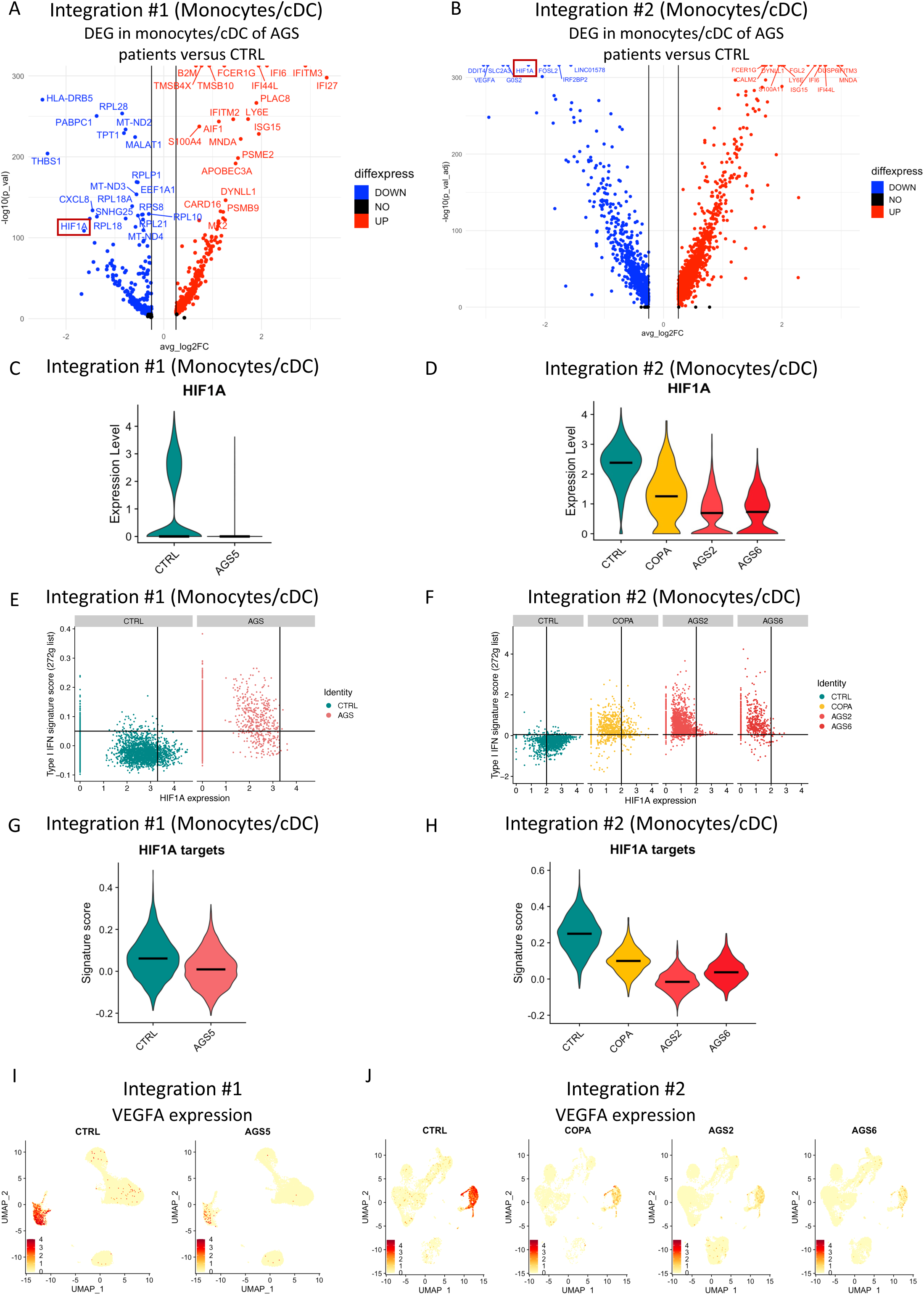
HIF1A expression and transcriptional activity are defective in monocytes/cDC of AGS patients. **(A and B)** Volcano plot of DEG in monocytes/cDC from AGS patients (AGS5) compared to CTRL of integration#1 **(A)** or from AGS patients (AGS2 + AGS6) compared to CTRL of integration#2 **(B)**. Top40 dysregulated genes are labelled in integration#1 and top20 dysregulated genes are labelled in integration#2, upregulated in red and downregulated in blue. DEG are displayed according to average log2 fold change (avg_log2FC) versus -log10 of adjusted *p* value (-log10(p-val)). Genes were considered modulated if avg_log2FC>0.25 or <- 0.25 and adjusted *p* value < 0.05. (C and D) Violin plot of *HIF1A* expression in monocytes/cDC split by CTRL, AGS2, AGS5 AGS6 and COPA groups from integration#1 (C) and from integration#2 (D). Medians are shown as black lines. (E and F) *HIF1A* expression versus type I IFN signature (272genes list) in monocytes/cDC split by CTRL, AGS2, AGS5 AGS6 and COPA groups from integration#1 (E) and from integration#2 (F). (G and H) Violin plot of the signature score of HIF-1α known targets expression from NCI- Nature database (“HIF-1-alpha transcription factor network Homo sapiens”) in monocytes/cDC split by CTRL, AGS2, AGS5, AGS6 and COPA groups from integration#1 (G) and from integration#2 (H). Medians are shown as black lines. (I and J) Feature plot of VEGFA expression in all PBMC split by CTRL, AGS2, AGS5, AGS6 and COPA groups in integration#1 (I) and integration#2 (J). In integration #1, Monocytes/cDC refers to clusters 5, 14, 17, 8, 20, 15 at the resolution 1.6 and in integration #2, Monocytes/cDC refers to clusters 5, 17, 23, 25, 27 at the resolution 1.6. See also figS3.

### Energy metabolism switch with mitochondrial stress in monocytes/cDC of AGS patients

To further explore the transcriptomic dysregulations observed in our scRNA-seq datasets, we assessed DEG between AGS and CTRL cells among monocytes/cDC to reveal cellular pathways potentially affected by the disease. We applied separately upregulated and downregulated DEG lists to several independent databases for pathway enrichment using EnrichR software ^23^. As expected, in both integrations, the top upregulated genes belonged to pathways directly related to type I IFN signaling (**fig4A and B**). However, we also observed an upregulation of metabolic pathways, in particular oxidative phosphorylation, that has not been previously reported in this context. Importantly, this observation was associated with a downregulation of genes associated with anaerobic glycolysis (also known as the Warburg effect) and HIF-1α signaling. We observed the same pathways when analysing DEG between AGS patients and CTRL among all PBMC (**figS4A and B**). Oxidative phosphorylation represents the most efficient source of energy from glucose, and involves the respiratory chain in mitochondria. In contrast, anaerobic glycolysis is less efficient in energy production, but more responsive, and does not require mitochondria oxygen consumption ^24^. HIF-1α stands at the fork between oxidative phosphorylation and anaerobic glycolysis, favouring glucose catabolism through the later and inhibiting oxidative phosphorylation. We further assessed energy metabolism in AGS and COPA patient cells by applying publicly available signatures for anaerobic glycolysis and oxidative phosphorylation pathways ^21, 25^. Accordingly, we observed that cells of AGS patients demonstrated increased expression of the genes involved in oxidative phosphorylation, together with a decreased in the expression of the genes involved in anaerobic glycolysis (**fig4C and D**). Interestingly, COPA patients’ cells exhibited, here again, intermediate signature scores for both metabolic pathways (**fig4D**), correlated with *HIF1A* expression and activity. When compared to other cell types, monocytes/cDC had the most noticeable increase in oxidative phosphorylation signature associated with a strong decrease in genes related to anaerobic glycolysis in patients (**figS4C and D**). Elevated expression of genes involved in oxidative phosphorylation led us to assess features of mitochondrial stress in transcriptomic data from patients. We applied a mitochondrial stress signature (**see supplementary information table SI2** ^26–31)^ to our dataset and observed a higher expression of mitochondrial stress genes in all PBMC of AGS patients and in particular in monocytes/cDC (**fig4E and F**). Again, this mitochondrial stress signature was intermediate in COPA patients’ cells (**fig4F**). Thus, the major transcriptomic changes observed in monocytes/cDC from patients with AGS, besides type I IFN response, reflect an energy metabolism switch towards oxidative phosphorylation associated with mitochondrial stress in AGS patients. Regarding this energy metabolic swithc, COPA patients presented an intermediate phenotype, correlated with an intermediate response to type I IFN and *HIF1A* expression.

**Fig 4:**
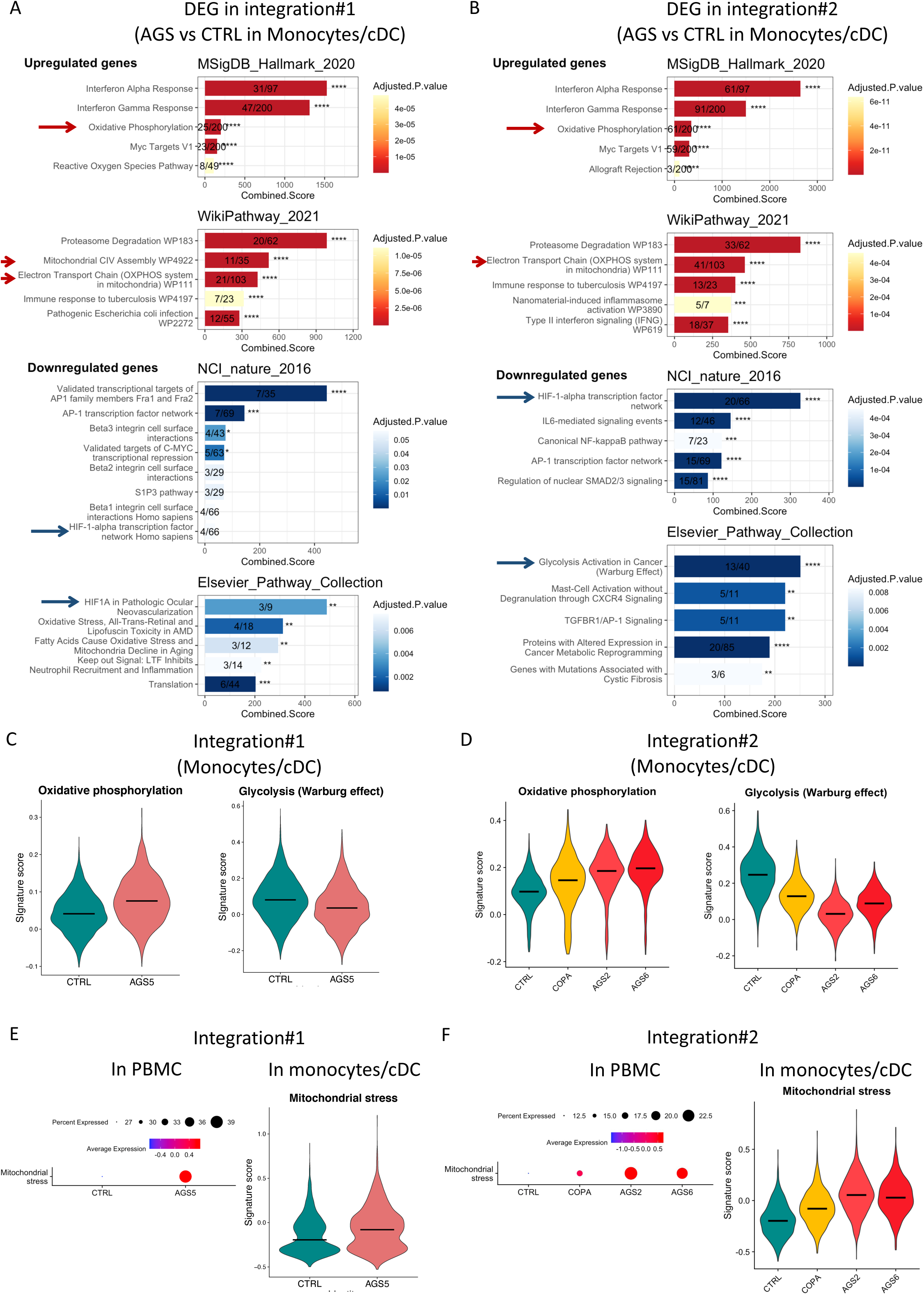
Energy metabolism switch in monocytes/cDC of AGS patients. **(A and B)** Pathway enrichment analysis using EnrichR (Chen et al. 2013) of DEG in AGS patients (AGS5) compared to CTRL of integration#1 **(A)** or in AGS patients (AGS2 + AGS6) compared to CTRL of integration#2 **(B)**. Pathways are ranked based on combined score. Color gradient is proportional to adjusted *p* value of Fisher’s test and labelled as ***:*p*<0.001, **:*p*<0.01, *:*p*<0.05. Numbers indicates the number of genes present in the DEG list over the total number of genes referenced in the corresponding pathway. Red highlight pathways including genes upregulated in patients and blue highlight pathways including downregulated genes in patients. Red arrows point toward Oxidative phosphorylation associated pathways and blue arrows point toward anaerobic glycolysis and *HIF1A*-related pathways. (C and D) Violin plot representing signature score of the gene list in “Oxydative phosphorylation pathway” (MSIgDB hallmark datatbase) (left panel) and of “Glycolysis Activation in Cancer Warburg Effect” (Elsevier database) (right panel) on monocytes/cDC split by CTRL, AGS2, AGS5, AGS6 and COPA groups from integration#1 (C) and integration#2 (D). Medians are shown as black lines. **(E and F)** Dotplot of “Mitochondrial stress” signature score in PBMC (left panel) and violin plot of “Mitochondrial stress” signature score in monocytes/cDC (right panel) split by CTRL, AGS2, AGS5, AGS6 and COPA groups from integration#1 **(E)** and integration#2 **(F)**. In dotplot, size of dots indicates percentage of cells in which the gene is detected while color indicate average expression of the signature. In violin plot, medians are shown as black lines. In integration #1, Monocytes/cDC refers to clusters 5, 14, 17, 8, 20, 15 at the resolution 1.6 and in integration #2, Monocytes/cDC refers to clusters 5, 17, 23, 25, 27 at the resolution 1.6. See also figS4.

### Detection by scRNA-seq of a monocyte/cDC cluster of cells specific to the response to the treatment with reverse transcriptase inhibitors (RTI)

As scRNA-seq allowed us to establish important transcriptomic changes in AGS patients particularly in monocytes/cDC, we aimed at following the transcriptomic profile of patients before and after an experimental treatment with reverse transcriptase inhibitors (RTI) triple therapy ^32^. We therefore focused on P2 harbouring biallelic *SAMHD1* mutations from whom we were able to generate scRNA-seq data before (P2_before_T), and 1 month after the initiation of treatment (P2_after_T) from integration#1. Both samples were integrated with data from controls C1 and C3 to generate integration#3 (**fig5A and S5A**). First, we assessed cell type proportions in each sample and observed a decrease in the proportion of memory T cells and NK cells and an increase in the proportion of B cells in patient#2 compared to the CTRL group (**figS5B**), as already described in integration#1 and #2 (**fig1E and 1F**). Then, we assessed the type I IFN response and confirmed that monocytes/cDC presented the greater score of type I IFN signaling (**figS5C**). As expected monocytes/cDC from patient before treatment (P2_before T) present a high type I IFN response compared to CTRL. Interstingly, after treatment (P2_after_T), we detected a portion of cells with a decreased type I IFN response (**fig5B**). Based on in silico clustering, these cells with a low type I IFN response gathered together in a unique cluster labelled cluster 12 (**fig5C**). This cluster was seen exclusively in the treated patient (**fig5D**). It consists in a mixture of cDC, classical monocytes and non-classical monocytes based on markers expression (respectively, *FCER1A*, *S100A8* and *FCGR3A*) (**figS5D**). We hypothesized that cluster 12 might correspond to treatment related cells of the monocytic lineage. We then further characterized cells belonging to this cluster, and detected higher levels of *HIF1A* expression compared to other patient cells, reaching CTRL levels of expression (**fig5E and F**). Consistant with previuous results (**fig3**), cells with restored *HIF1A* expression demonstrated very low levels of type I IFN signature (**fig5G and S5E**). The activity of HIF-1α was also restored in cells of cluster 12, based on the elevated expression of HIF-1α targets in the treatment related cluster (**fig5H**). By performing DEG analysis between cells cluster 12 and other monocytes/DCs of P2 after treatment, we confirmed a downregulation of genes belonging to type I IFN response and to oxidative phosphorylation (**fig5I and J**). Indeed, cells from cluster 12 had a lower level of oxidative phosphorylation signature associated with a lower mitochondrial stress score, compared to monocytes/cDC from P2, reaching a similar profile to the CTRL group (**fig5K, 5L**). Those results suggest that the treatment impacted a fraction of patients’ monocytes/cDC with a reduced type I IFN response alongside a restored HIF-1α activity, reduced oxidative phosphorylation and decreased mitochondrial stress. Those results further reinforce the observed link between the type I IFN response and HIF-1α regulation of energy metabolism in AGS.

**Fig 5:**
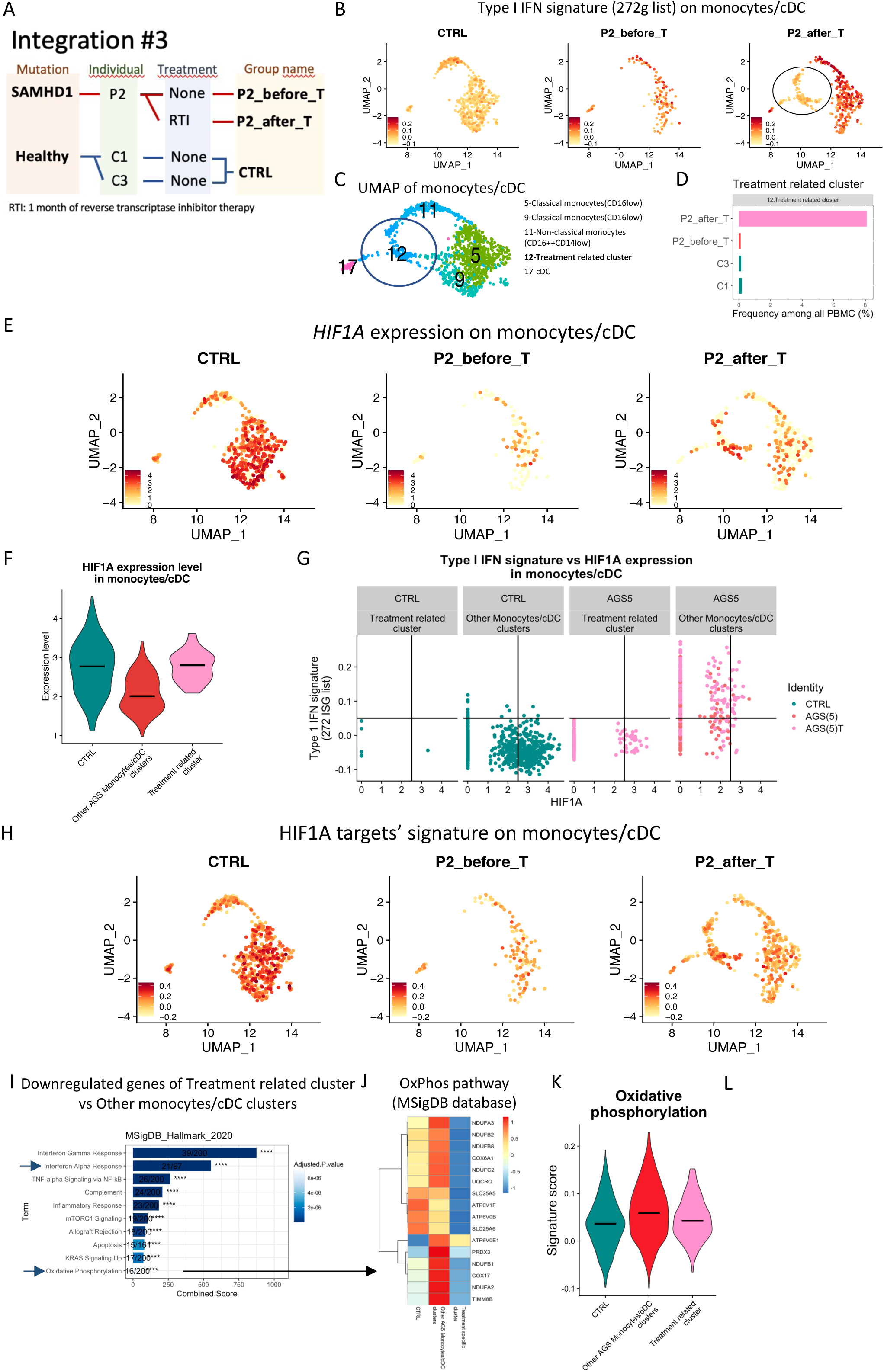
Detection of treatment-related cells by scRNA-seq in one AGS patient. (A) Integration of PBMC analyzed at the scRNA-seq level form P2 before (P2_before_T) and after 1 month of treatment (P2_after_T) with RTI tri-therapy. P2 is mutated on *SAMHD1* genes and developed AGS. During the experiment, PBMC form two unrelated healthy individuals (C1 and C3) were added and integrated to generate integration#3. (B) Feature plot of type I IFN signature score (272 ISG list) focused on monocytes/cDC and split by CTRL, P2_before_T and P2_after_T groups. Blue circle indicates a cluster unique to P2_after_T patient with low type I IFN signature score. (C) Cluster identification of monocytes/cDC in integration#3 at the 1.6 resolution. Blue circle indicates the cluster 12, labelled “treatment related cluster”. (D) Frequency of cell in the treatment related cluster among PBMC from each sample of integration#3. (E) Feature plot of *HIF1A* expression in monocytes/cDC split by CTRL, P2_before_T and P2_after_T groups of integration#3. (F) Violin plot of *HIF1A* expression in the treatment related cluster against other AGS monocytes/cDC clusters from P2_after_T and P2_before_T samples. Only *HIF1A* expressing monocytes/cDC are shown. Medians are shown as black lines. (G) *HIF1A* expression against ISG signature score (272 gene list) of monocytes/cDC in integration#3 split by cells belonging to the treatment related cluster or not. Each dot is a cell and green represent CTRL group while dark pink are cells belonging to P2_before_T and light pink cells belonging to P2_after_T. (H) Feature plot of the signature score of HIF-1α known targets expression from NCI- Nature database (“HIF-1-alpha transcription factor network Homo sapiens”) in monocytes/cDC split by CTRL, P2_before_T and P2_after_T groups from integration#3 (H). Medians are shown as black lines. **(I)** Pathway enrichment analysis made with EnrichR form downregulated genes in the treatment related cluster compared to other monocytes/cDC from the treated patient (P2_after_T). Pathways are ranked based on combined score. Color gradient is proportional to adjusted *p-*value of Fisher’s test and labelled as ***:*p*<0.001, **:*p*<0.01, *:*p*<0.05. Numbers indicates the number of genes present in the DEG list over the total number of genes referenced in the corresponding pathway. Arrows point toward type I IFN and oxidative phosphorylation associated pathways. (J) Heatmap of the gene from the oxidative phosphorylation apthway from MsigDB-Hallmark- 2020 database shown in I on CTRL, treatment related cluster and other AGS monocytes/cDC clusters groups. (K) Violin plot of “Oxidative phosphorylation pathway” (MSIgDB hallmark database) signature score in the treatment related cluster against CTRL group and other AGS monocytes/cDC clusters. Medians are shown as black lines. **(L)** Violin plot of “Mitochondrial stress” signature score in the treatment related cluster against CTRL group and other AGS monocytes/cDC clusters. Medians are shown as black lines. In integration#3, “Monocytes/cDC” refers to clusters 5, 9, 11, 12, 17 at 1.6 resolution. In fig5, clusters are grouped as follows: “Treatment related cluster” (cluster 12 from P2_after_T), “Other AGS Monocytes/cDC clusters” (clusters 5, 9, 11, 17 from both P2_before_T and P2_after_T samples). See also figS5.

### Validation of metabolic switch and mitochondrial stress associated with a type I interferon response in an *in vitro* cellular model of AGS

To validate the observations drawn from scRNA-seq analysis of AGS patients’ material, we established an *in vitro* cellular model, based on primary monocytes from healthy blood donors that were differentiated into dendritic cells (MDDC, Monocyte Derived Dendritic Cell). In this model, we targeted *SAMHD1* expression at the mRNA level with short hairpin RNA (shRNA) to replicate its loss of function in AGS5 patients. We obtained 2 shRNA (shSAMHD1_3 and shSAMHD1), both efficiently inhibiting *SAMHD1* at the mRNA level (**fig6A and S6A**), with shSAMHD1 also presenting a noticeable downregulation at the protein level (**fig6B**). Upon knock-down (KD) of *SAMHD1*, MDDC produced detectable amounts of IFN-β protein and high amounts of IP-10 but not IL-1β or TNFα (**fig6C and S6B**), consistent with what has been reported in AGS patients. ISG15 expression, a hallmark of a type I IFN-mediated response, was tracked by flow cytometry staining (**figS6C**). ISG15 protein expression was significantly upregulated in shSAMHD1 transduced MDDC (**fig6D, 6E and S6D**), reaching a maximal response at day 6 after transduction (**figS6E**). A type I IFN response was also observed at the mRNA level by measuring the representative six-ISG signature described above (comprising: *ISG15, IFI44L, IFIT1, IFI27, RSAD2* and *SIGLEC1*) (**fig6F and S6F**). We aimed at confirming our observations related to *HIF1A* dysregulation in AGS, and detected lower amounts of HIF- 1α protein in SAMHD1-deficient MDDC (**fig6G**). We then performed Seahorse XF Analyser (Agilent) experiment to address potential energy metabolism imbalance resulting from observed HIF-1α dysregulation. Briefly, at day 4 after transduction with shRNA, extracellular acidification rate (ECAR), which is a readout of the lactate secretion by cells, was monitored to measure anaerobic glycolysis activity. In parallel, oxygen consumption rate (OCR) was measured reflecting oxidative phosphorylation activity through oxygen consumption by the complex 5 of the mitochondrial electron transport chain. Those measurements also allows to quantify ATP synthase activity, maximal respiratory capacity, actual glycolysis and maximal glycolytic capacity by adding a panel of drugs during the time course (**see supplementary information SI3**). These experiments revealed that shSAMHD1-transduced MDDC had reduced glycolytic capacity (**fig6H**). Reduced glycolytic capacity was associated with qRT- PCR results showing increased expression of the mitochondrial stress associated genes *CLPP* and *HSP60* in SAMHD1-deficient MDDC (**fig6J**). All these results were further validated when using another shRNA targeting SAMHD1 (shSAMHD1_3) (**figS6A to F**). Finally, we used a cocktail of type I IFN-blocking antibodies and observed that ISG15 expression and IP-10 production in SAMHD1-deficient MDDC were lost, confirming the strict type I IFN dependency of these cellular responses (**fig6K, 6L and S6G**), in our *in vitro* system. Here, by generating an *in vitro* cellular model of AGS5 in human primary MDDC, we reproduced spontaneous type I IFN production followed by response to type I IFN, as well as high IP-10 secretion, alongside reduction of HIF-1α expression and dampened anaerobic glycolysis. This model functionally validates what was observed at the scRNA-seq level in monocytes/cDC of AGS patients.

**Fig 6:**
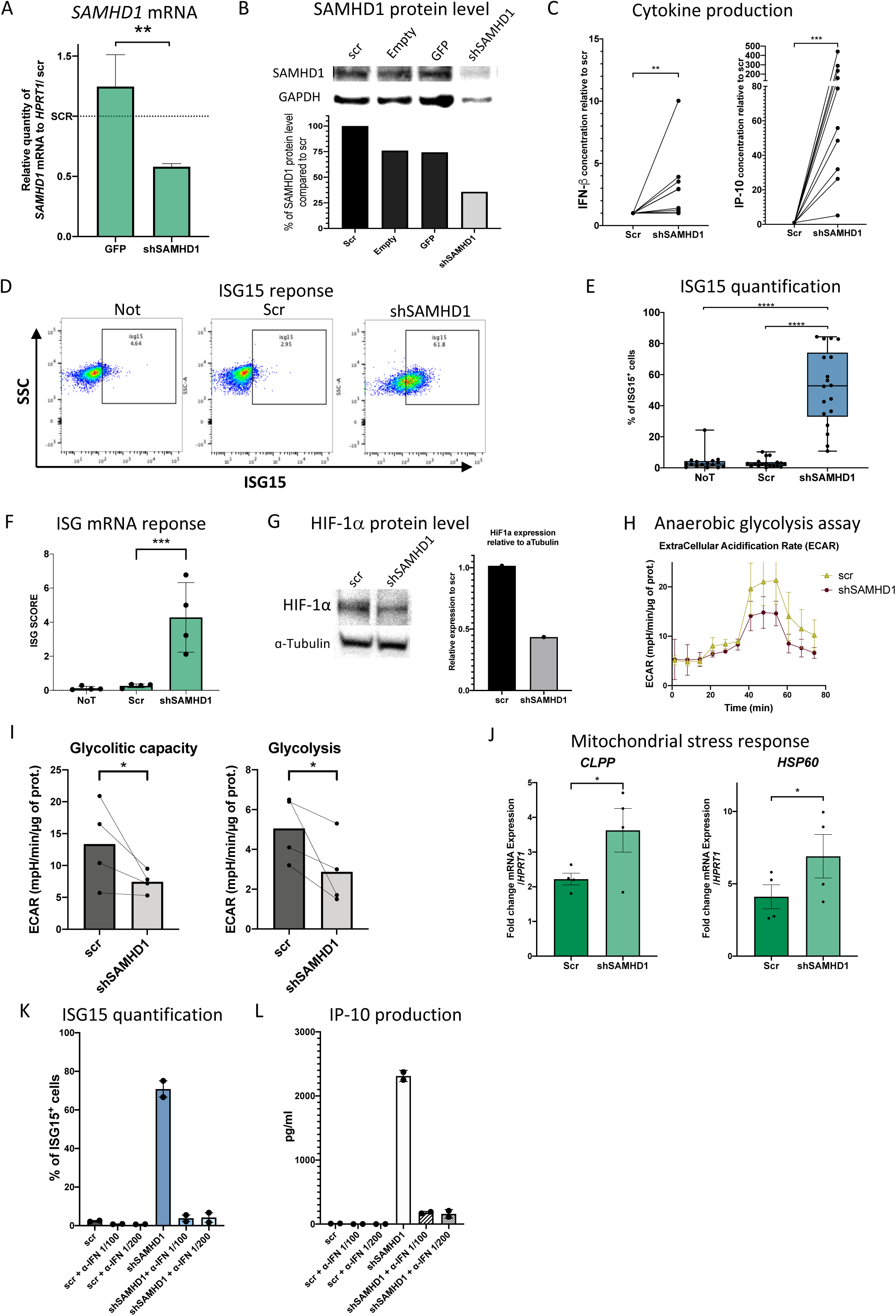
Validation of the metabolic switch, mitochondrial stress and type I inteferon response in an *in vitro* cellular model of AGS. MDDC differentiated from monocytes of healthy donors over 4 days and transduced by shRNA targeting *SAMHD1* mRNA (shSAMHD1) as soon as day 0. Two shRNA were selected as potent SAMHD1 inhibitors, one is presented in main fig6 and the other in figS6. **(A)** RT-qPCR of *SAMHD1* on RNA extracted at day 5 from MDDC transduced by lentiviruses bearing an empty vector (Empty) or coding for scr, GFP, or shSAMHD1 on 3 unrelated healthy donors. Data have been normalized to *HPRT1* gene expression and represented as fold change to scr condition. Anova statistical test is represented as: ***:*p*<0.001, **:*p*<0.01, *:*p*<0.05. **(B)** Western blot of SAMHD1 protein level in transduced MDDC. Proteins were extracted at day 6 and revealed for SAMHD1 and GAPDH (Glyceraldehyde-3-phosphate dehydrogenase) expression (top panel). Bar graph of quantified western blot band for SAMHD1 normalized to GAPDH protein level and represented as percentage of the level obtained in scr condition. **(C)** Cytokine measurement by legendplex in supernatant of MDDC at day 6 post transduction coming from 11 unrelated healthy donors. Data are represented as fold change to scr condition. Wilcoxon’s statistical test is represented as: ns: not significant, ***:*p*<0.001, **:*p*<0.01, *:*p*<0.05. (D) Representative flow plots showing intracellular staining of ISG15 plotted versus SSC (side scatter) parameter of MDDC at day 6 post transduction. Gating was positioned according to ISG15 isotype control staining, shown in figS6D. **(E)** Quantification of the percentage of ISG15 positive cells in flow cytometry among MDDC at day 6 post transduction. Data coming from MDDC generated from 18 unrelated healthy donors. (F) ISG score based on mRNA quantification by RT-qPCR of *ISG15, IFI44L, IFIT1, IFI27, RSAD2* and *SIGLEC1* in MDDC at day 5 after transduction from 4 unrelated healthy donors. Score refers to the mean of relative mRNA level normalized to *HPRT1* of all 6 genes. (G) Western blot of HIF-1α protein level from protein extracted from MDDC, 5 days after transduction and revealed for HIF-1α and α-Tubulin (left panel). Quantification of HIF-1α protein level relative to α-Tubulin level is shown represented as fold change to scr condition (right panel). (H) Representative time course of Seahorse XF Analyzer experiment from MDDC at day 6 after transduction. Extracellular acidification rate (ECAR) was measured and normalized to the quantity of protein in each well. Mean with standard deviation of 3 replicates from 1 healthy donor are represented. (I) Quantification of extracellular acidification rate (ECAR) due to glycolytic capacity or actual glycolysis from 4 unrelated healthy donors. **(J)** RT-qPCR of *CLPP* and *HSP60* genes on RNA extracted from MDDC at day 5 after transduction. Data coming from 4 unrelated healthy donors are represented normalized to *HPRT1* mRNA level. (K and L) At day 4 after transduction, increasing amount of type I IFN blocking antibodies cocktail (α-IFN) (1/100 and 1/200 dilution factor) were added to MDDC culture for 48 hours. At day 6 post transduction cell were stained for intracellular ISG15 expression by flow cytometry (K) and supernatant was harvested for quantification of IP-10 production by legendplex (L). (D, E and F) Mean with SEM (standard error of the mean) are represented with one way anova with Sidak multiple correction statistical test as ns: not significant, ***:*p*<0.001, **:*p*<0.01, *:*p*<0.05. **(I and J)** Mean and SEM are represented with paired t-test statistics as: ns: not significant, ***:*p*<0.001, **:*p*<0.01, *:*p*<0.05. (A to L) NoT: Not transduced MDDC; Empty: MDDC transduced with an empty vector; scr: MDDC transduced with a scramble RNA without known target in human cells; GFP: MDDC transduced with a GFP coding plasmid for assessment of transduction efficiency; shSAMHD1: MDDC transduced with the most efficient shRNA against *SAMHD1*. See also figS6.

### Stabilization of HIF-1α expression level reverts the energy metabolism switch, relieves mitochondrial stress and inhibits IP-10 production

As HIF-1α activity is known to control energy metabolism, we investigated whether we could correct the metabolic switch observed, in our *in vitro* model of SAMHD1-deficient MDDCs by using the prolyl hydroxylase inhibitor dimethyloxalylglycine (DMOG), known to prevent HIF- 1α degradation. 500 µM of DMOG effectively increased HIF-1α protein levels as early as 5 hours post incubation, although less effectively in SAMHD1 KD cells due to decreased HIF- 1α level in these cells (**fig7A**). To monitor HIF-1α function, the energy metabolism state of SAMHD1-deficient MDDC was assessed by Seahorse XF Analyser (Agilent), in the presence or absence of two different doses of DMOG (250 µM and 500 µM) for 48 hours. We noted that HIF-1α stabilisation reduced the oxidative phosphorylation capacity of cells, while anaerobic glycolysis activity was increased, in a dose dependent manner (**fig7B**). Taken together, these results suggest that glucose consumption was redirected from mitochondrial respiration toward anaerobic glycolysis. Consistently, following DMOG treatment, *CLPP* and *Hsp60* gene expressions were reduced in SAMHD1-deficient MDDC, indicating a reduced mitochondrial stress (**fig 7C**). We then assessed the impact of DMOG on type I IFN production and response in our model. HIF-1α stabilisation partially reduced mRNA levels of some ISG (**fig7D and S7C**), but did not affect ISG15 protein levels (**fig7E and S7D**), nor IFN-β production (**fig7F**). However, SAMHD1-deficient MDDC produced between 2 to 4 times lower amount of IP-10 following DMOG treatment (**fig7F**). Similar results were reproduced with shSAMHD1_3 (**figS7A to E**). In conclusion, we are reporting that HIF-1α stabilisation by DMOG reverts the energy metabolism switch observed in SAMHD1-deficient MDDC, relieves mitochondrial stress and markedly inhibits the production of the neurotoxic cytokine IP-10.

**Fig 7:**
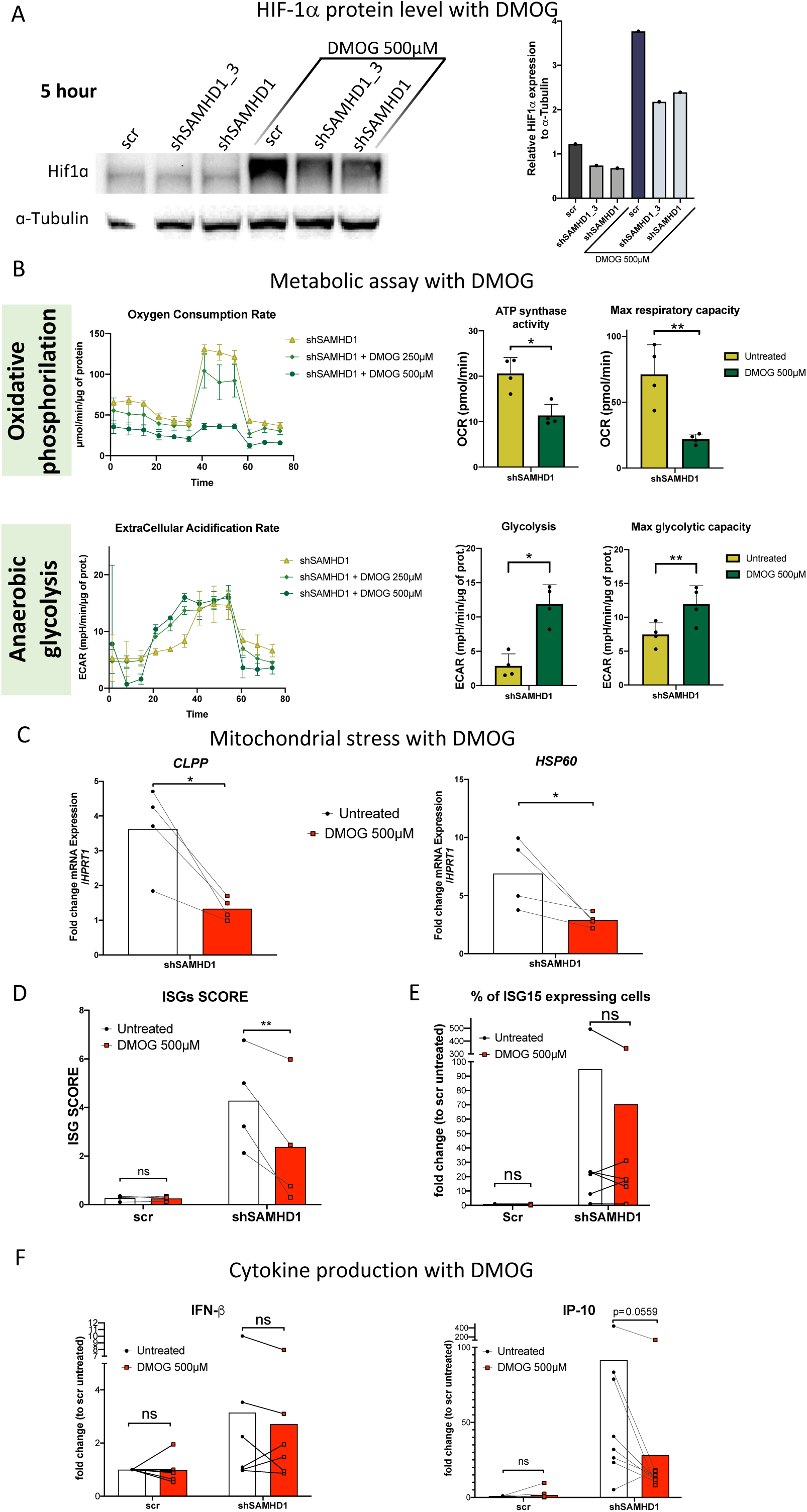
Stabilization of HIF-1ɑ, at the protein level, reverts energy metabolism switch, releases mitochondrial stress and inhibits IP-10 production. At day 4 after transduction, MDDC transduced with shRNA targeting *SAMHD1* were treated with the HIF-1ɑ stabilizing drug dimethyloxalylglycine (DMOG) for indicated time and concentration. DMOG is an antagonist of the HIF prolylhydroxylase cofactor that is necessary for HIF-1ɑ degradation by the proteasome. (A) MDDC at day 4 after transduction were treated with 500 µM of DMOG for 5 hours before being lysed for protein extraction. HIF-1ɑ level of protein expression observed by western blot revelation is represented and compared to ɑ-tubulin (left panel). Quantification of HIF-1ɑ protein level is normalized to ɑ-tubulin protein level (right panel). (B) MDDC at day 4 after transduction were treated with increasing concentration of DMOG (250 µM and 500 µM) for 48 hours before undergoing the Seahorse XF Analyzer time course to determine oxygen consumption rate (OCR) (top panels) and extracellular acidification rate (ECAR) (Bottom panels). A Seahorse XF Analyzer time course is represented for 1 donor for both assays showing mean measurement with standard error of triplicates at each time point (Left panels). ATP synthase activity, Max respiratory capacity, actual glycolysis and max glycolytic capacity were quantified for 4 unrelated healthy donors assessed by Seahorse XF Analyzer experiment untreated or with 500 µM of DMOG (right panels). **(C)** RT-qPCR of *CLPP* and *HSP60* on RNA extracted from MDDC at day 5 after transduction (24 hours of DMOG treatment) for 4 unrelated healthy donors. Data were normalized to the level of expression of the *HPRT1* gene measured in each sample. (D) ISG score based on mRNA quantification by RT-qPCR of *ISG15, IFI44L, IFIT1, IFI27, RSAD2* and *SIGLEC1* genes on MDDC at day 5 post transduction (24 hours of DMOG) from 4 unrelated healthy donors. Score refers to the mean of relative mRNA level normalized to *HPRT1* of all 6 genes. (E) Quantification of the percentage of ISG15 positive cells in flow cytometry among MDDC at day 6 post transduction (48 hours of DMOG) from 6 unrelated healthy donors. **(F)** Cytokine measurement by legendplex in supernatant of MDDC at day 6 post transduction (48 hours of DMOG) for 6 unrelated healthy donors. Data are represented as fold change to scr condition. **(B and C)** Means and paired T test statistics are represented as: ns: not significant, ***:*p*<0.001, **:*p*<0.01, *:*p*<0.05 **(D, E and F)** Mean and one way anova with Sidak multiple correction statistical test are represented as: ns: not significant, ***:*p*<0.001, **:*p*<0.01, *:*p*<0.05. See also figS7.

## Discussion

### Characterization of AGS pathogenesis at the molecular level

Our findings from scRNA-seq analysis of PBMC from AGS patients, validated in our *in vitro* cellular model, are summurized in the following conceptual model, in accordance with existing literature (figS7E, F and G). In AGS, our results revealed a metabolic switch toward oxidative phosphorylation with reduced anaerobic glycolysis orchestrated by a loss of HIF-1α expression and activity. The results further suggest that constant type I IFN production induces molecular processes that inhibit HIF-1α stabilisation through reduced mRNA expression (**figS7F**). Oxidative phosphorylation is the favoured pathway to generate energy for being the most efficient way to process glucose but this metabolic pathway has the downside of being a major source of ROS, a known inducer of oxidative stress. In healthy individual, HIF-1α, a sensor of oxidative stress, is stabilized at the protein level upon ROS accumulation, and regulates ROS production by favouring glucose processing through anaerobic glycolysis, bypassing oxidative phosphorylation (**figS7E**). In AGS patients’ cells, HIF-1α expression is supressed. Thus, ROS can accumulate, increasing oxidative stress and may induce several events such as NF-κB activation and mitochondrial stress. While NF-κB activation is known to aggravate an IP-10 response, chronic mitochondrial stress affects mitochondrial membrane permeability, favouring mtDNA/RNA release in the cytoplasm. As cytoplasmic nucleic sensing pathways can detect mtDNA/RNA, we propose that this feeds a vicious cycle of type I IFN production and chronic IP-10 production in AGS where nucleic acid sensing pathways are already sensitized by genetic mutations (**figS7F**). This mechanism is expected to be even more important in cells with high ROS content such as myeloid cells. In an *in vitro* cellular model of AGS, HIF-1α stabilisation by the synthetic chemical compound DMOG, allowed a switch of energy metabolism to anaerobic glycolysis, even in presence of type I IFN. In this way, mitochondrial stress is relieved and IP-10 production is reduced (**figS7G**).

### Role of HIF-1α in monocytes/cDC

*HIF1A* downregulation was associated with a decreased expression of its target genes, especially in monocytes/cDC of AGS patients. HIF-1α is a sensor of metabolic stress in cells, and is stabilized when ROS reach a dangerous levels or if oxygen is lacking ^18^. Several sources of ROS can be listed, with the main one involving electron leakage from the mitochondria electron transport chain ^33^. HIF-1α stabilisation inhibits mitochondrial activity in order to supress ROS generation and oxygen consumption by the electron transport chain complexes ^19^. To compensate for the lack of energy production following mitochondria inhibition, conversion of glucose into lactate (anaerobic glycolysis) is favoured. Several enzymes of the anaerobic glycolysis are upregulated by HIF-1α, including the lactate dehydrogenase that transforms the glycolysis end product pyruvate into lactate ^19^. The switch of energy production from mitochondria activity toward anaerobic glycolysis is called the Warburg effect, first described in cancer cells ^34^. Later studies have established the importance of this energy metabolism switch in critical cellular processes such as immune cell proliferation ^35^. We hypothesize that *HIF1A* is more highly expressed in monocytes/cDC, as these cells rely more heavily on the Warburg effect than other cell types. Indeed, upregulation of anaerobic glycolysis has been shown to be necessary for the proper activation and maturation of dendritic cells ^36^ and macrophages ^37^. On top of that, monocytes rely on ROS generation for pathogen clearance ^38^, and dendritic cells antigen presentation capacity also involve ROS ^39^, implying a greater susceptibility to ROS induced stress and, consequently, HIF-1α stabilisation. Our results place *HIF1A* at the centre of the regulation of monocytes/cDC at the transcriptomic level, in accordance with current literature establishing the importance of energy metabolism regulation in those cells. Altogether, scRNA-seq data in AGS patients suggest a close relationship between elevated type I IFN response and low levels of *HIF1A* expression. To our knowledge, only one study reported a potential link between type I IFN and HIF-1α, showing ISG15 capacity to decrease HIF-1α protein levels through proteasome induced degradation without affecting mRNA levels ^20^. These results could indicate that a chronic type I IFN exposure, like in AGS, leads to *HIF1A* inhibition at the mRNA level. Our results on COPA patients further support a close regulation of *HIF1A* by type I IFN response. Like AGS, COPA is a monogenic disease associated with enhanced type I IFN signaling ^13^ but differs in clinical manifestations ^13^. scRNA-seq analysis revealed that COPA patients’ cells exhibit an intermediate level of type I IFN response which was associated with an intermediate level of *HIF1A* inhibition compared to AGS patients’ cells. While ISG15 likely has a role, we do not exclude potential role of other ISG in the regulation of *HIF1A*. Finally, type I IFN have been shown to favour genes involved in oxidative phosphorylation over anaerobic glycolysis in human monocyte-derived cells ^40^, coherent with a reduce *HIF1A activity*.

### Uncontrolled oxidative phosphorylation induces mitochondrial stress

We observed an increased expression of genes involved in oxidative phosphorylation in AGS patients’ cells associated with decreased signatures of anaerobic glycolysis, especially in monocytes/cDC. Moreover, cells from AGS patients presented increased expression of mitochondrial stress genes. Altogether, these data suggest that lack of *HIF1A* expression confers a defect in cell capacity to regulate oxidative phosphorylation by switching to anaerobic glycolysis. We hypothesize that overactivation of oxidative phosphorylation, both in intensity and in time, favours mitochondrial stress. Two independent teams recently performed genome wide association studies (GWAS) on large cohorts to identify single nucleotide polymorphisms (SNP) associated with disruption in mtDNA copy number ^41, 42^. Both studies revealed that SNP in *SAMHD1* correlate with an increase in mtDNA copy number, strengthening our hypothesis of high mitochondrial stress in AGS (at least in *SAMHD1* mutated patients) as disruption of mtDNA copy number is indicative of oxidative stress in the mitochondria ^43, 44^. One good candidate for oxidative phosphorylation induction of mitochondrial stress is ROS generation by the electron transport chain complexes. ROS accumulation has been shown to be an inducer of mitochondrial stress which, if uncontrolled, leads to a loss of mitochondria integrity ^45^. Thus, we speculate that lack of *HIF1A* expression in AGS would result in defective sensing of ROS accumulation from oxidative phosphorylation, in turn inducing mitochondrial stress. Accordingly, when HIF-1εξ was stabilized in our *in vitro* cellular model of SAMHD1 deficiency, oxidative phosphorylation was reduced, and mitochondrial stress relieved. Of possible relevance, we note that AGS has been suggested to involve a disturbance of mitochondria activity in some patients ^46, 47^. Notably, a recent report revealed alteration of mitochondrial integrity associated with uncontrolled oxidative phosphorylation, increased ROS activity and mitochondrial stress in AGS patients mutated in *RNASEH2A* or *RNASEH2B* ^48^. We propose here, that HIF-1εξ loss of activity could be a driver of mitochondrial dysfunction in AGS patients. It has been shown that chronic mitochondrial stress drives a loss of mitochondrial membrane integrity ^45^. Consequently, mtDNA and mtRNA can be released into the cytoplasm and then sensed by nucleic acid receptors, resulting in type I IFN production ^45, 49, 50^.Thus, it is tempting to speculate that type I IFN inhibition of *HIF1A* expression could further fuel nucleic acid sensing in AGS through uncontrolled mitochondrial activity followed by mtDNA/RNA release in the cytosol, resulting in exacerbated inflammation.

### Uncontrolled production of IP-10 in AGS, partly driven by metabolic switch

We also observed high IP-10 production in MDDC KD for SAMHD1. IP-10 is one of the most highly expressed inflammatory cytokines detected in plasma and cerebrospinal fluid (CSF) of AGS patients ^10^. Its proinflammatory role has been attributed to an induction of chemotaxis of inflammatory cells bearing the CXCR3 receptor, which includes monocytes, NK cells, activated T cells and neutrophils ^51^. Of note, IP-10 can also negatively regulate angiogenesis ^52^ which might relate to the high frequency of vasculitic chilblain-like lesion seen in patients with AGS. Moreover, IP-10 possess neurotoxicity capacity ^11^, which might be correlated with deleterious neurological manifestations in AGS. Factors disturbing brain development in AGS are unclear and most inflammatory cytokine have been shown to have deleterious effect on brain development in the context of viral infection during pregnancy, including IL-6, IL-1β, IL- 17A or IFN-γ ^53^. However, the particular context of AGS involves a relatively low production of IL-8 and IL-6 and no expression of IL-17A or IFN-γ in CNS of patients compared to viral infection ^10^. Another cytokine found elevated in AGS and having neurotoxic capacity is CCL2, mainly through recruitment of inflammatory cells, but were not assessed in our study ^54^. We wanted to focus on initiator events and consequently focus our analysis on the IFN inducible cytokine IP-10 for its high prevalence in AGS patients. Cellular sources of IP-10 are uncertain in AGS, endothelial and epithelial cells are potent producers, but some studies suggest that immune cells represent important IP-10 producers in human neurological disease ^55^. We show that, among PBMC, cells with the highest score for type I IFN response are monocytes/cDC. Moreover, *in vitro,* MDDC deficient for SAMHD1 produced high levels of IP-10, arguing for an important role for monocytes/cDC in the production of IP-10 in AGS patients. Nonetheless, this IP-10 production was reduced following HIF-1α stabilisation by DMOG, indicative of a suppressive effect of HIF-1α on IP-10 production. It has been shown that regulation of energy metabolism, in particular by HIF-1α, is crucial for DC maturation and other cytokines production ^36, 56^. Thus, we hypothesize here that high levels of oxidative phosphorylation are required for IP-10 production in MDDC and that HIF-1α stabilisation limits IP-10 production by reverting energy metabolism towards anaerobic glycolysis. We show here that HIF-1α stabilisation inhibits IP-10 production without affecting type I IFN production, suggesting that high IP-10 secretion in AGS patient is favoured by HIF1A reduced expression in monocytes/cDC.

### Treatment related cluster of cells detected by scRNA-seq

Using scRNA-seq data we were able to detect cells responsive to RTI treatment in one patient with AGS. Responsive cells grouped in a unique cluster composed of subpopulations of both monocytes and cDC. These cells share a reduced type I IFN response compared to other monocytes/cDC correlated with restored *HIF1A* expression and reduced mitochondrial stress. A phase I clinical trial with RTI therapy over 12 months showed a reduced IFN-α levels in serum and CSF, and a lower type I IFN score in the whole blood of patients with AGS1-5 mutations ^32^. This indicates that circulating cells in treated patients are likely to be exposed to lower levels of type I IFN allowing some cells to escape ISG induction. Although, this result needs to be reproduced in other patients, it highlights the potential importance of performing single-cell analyses in finely monitoring the effect of a treatment at the molecular level, and its potential value as a new tool to be used in personalized medicine performed at the molecular level. This encourages future research studies to assess individual follow up of transcriptional profile by scRNA-seq during therapy, particularly in diseases with heterogeneous treatment responses.

### Limitations of the study

The main limitation in the use of scRNA-seq techniques is gene dropout, as it is estimated that only the most expressed genes are detected (around top 15-20% in each cell, depending on the cell type). Thus, the role of lowly expressed genes is overlooked. scRNA-seq technology is evolving rapidly, and in our study the detection levels have improved significantly while producing our different datasets, thus we preferred to not integrate together some produced data, in order to reduce potential batch effects. However, we were able to draw similar conclusions on AGS patients despite exploring independent experiments and integrations of scRNA-seq data from patients with different mutations, reinforcing our conclusions on AGS pathological mechanism. AGS is a rare genetic disorder, so we present here a study on a total of 6 patients of different genotypes. In our study, the limited number of samples and potential discrepancies are addressed through a combination of supervised (classical DEG) with unsupervised (GRN) approaches for scRNA-seq analysis and *in vitro* validation. We used the DMOG drug to stabilize HIF-1α in a more controlled context *in vitro* on a unique cell type population. While it proved to be helpful to better understand the importance of energy metabolism in AGS, we do not recommend it yet as potential therapeutic candidate. It would force the system into an anaerobic glycolytic state with potential harmful consequences. Indeed, while HIF-1α stabilizing drugs are in development and under clinical phase trials in the context of anaemia, safety concerns have been already raised relating to their long terms use ^57^. Despite this, our results, which highlight energy metabolism at the centre of AGS disease, are still suggesting that potential therapeutic approaches targeting mitochondrial stress, ROS accumulation or TCA cycle intermediates could be of interest. Another limitation of our study was the focus on circulating immune cells, our area of expertise, but we do not exclude that energy metabolism imbalance could be relevant in non-circulating cell types, and could have direct disruptive effect on cells such as neurones or glial cells during brain development.

Finally, we used scRNA-seq data to monitor the effect of treatment effectiveness in one SAMHD1 deficient patient treated with RTI. While the generated results are interesting, they require follow-up in a larger number of patients.

## Star Methods

### Key resources table

### Lead contact

Further information and requests for resources and reagents should be directed to and will be fulfilled by Mickaël Ménager

### Experimental model and subject details

#### Patients

Patients were aged between 3 and 16 years old with the exception of P8 being 36 years old and were included based on molecular diagnosis of mutations in *SAMHD1*, *RNASEH2B* or *ADAR1* genes for AGS or *COPA* gene for patients with COPA disease. P1, P2, P3 and P4 studied in this work have been previously reported by Rice at al. respectively labelled P0011, P0003, P0002 and P0004 ^32^. PBMC were frozen the day of sampling and thaw once for scRNA-seq library generation. P1 was undergoing RTI tri-therapy (zidovudine, abacavir, lamivudine) for 5 months. PBMC from P2 were extracted before and after 1 month of RTI tri-therapy (zidovudine, abacavir, lamivudine). Other patients and healthy donors were not under therapy. The study was approved by the *Comité de protection des personnes Iles de France II* and the French advisory committee on data processing in medical research (ID-RCB: 2014-A01017-40). Consent of parents and/or patients, depending on age, was obtained for conducting the experiment.

#### Cells

To generate monocytes derived dendritic cells (MDDCs) form human primary cells, we acquired buffy coats from healthy human donors (*Etablissement Français de Sang*). We performed Ficoll extraction of peripheral blood mononuclear cells (PBMC) and used anti-human CD14 magnetic beads (Miltenyi) to purify monocytes. Magnetic sorting was performed on AutoMACS pro separator device (Miltenyi Biotech). Isolated monocytes were cultured in RPMI (Gibco), 10% FBS (heat inactivated, Sigma), 10 mM Hepes, 55 µM β-mercaptoethanol, 6 mM L-glutamine, 50 µg/ml Gentamicin and 100 U/ml Penicilin/Streptomycin (Gibco) in the presence of 10 ng/mL of human recombinant GM-CSF and 50 ng/mL of human recombinant IL-4 (Miltenyi) to induce MDDC differentiation ^58^. Multiple lot of FBS were tested and selected according minimal induction of CD86 expression on MDDC. 5x10^6^ cells/mL of CD14^+^ cells were plated in 10 cm culture plates (Sigma) and cultured for 4 days before functional assays to allow complete MDDC differentiation which was assessed by flow cytometry following CD14 loss of expression and DC-SIGN upregulation. Fresh media with cytokines (20% of total volume) was added at day 1 and day 3. At day 4 of cultures, cells were washed with fresh medium, counted and plated on 96 wells culture plates (Falcon) for stimulation and functional assays at the indicated times.

PBMC collected from patients were isolated by Ficoll-Paque density gradient (Lymphoprep, Proteogenix) from blood samples using standard procedures. PBMC were frozen in liquid nitrogen in DMSO 10% FBS 90%. scRNA-seq experiment were performed immediately after thawing.

### Method details

#### Plasmids and viral particle production

SAMHD1 shRNA coding plasmids were purchased from SIGMA company and are detailed in key resources table. Vpx coding plasmid was used to allow monocytes transduction as previously described ^59^. pLK0.1-GFP was used to assess transduction efficiency while pCMV- ΔR8.91 and pCMV-VSVg were used for packaging and pseudotyping of viral particles as previously described ^60^. Plasmid amplification was achieved by transformation of Stblt3 bacteria (ThermoFisher) and ampicillin selection. All plasmids were then purified using Purelink Hipure plasmid midiprep kit (ThermoFisher). To produce VLP we transfected plasmids into HEK293FT cells as previously described ^61^. HEK293FT cells (Invitrogen) were cultured in DMEM (Gibco), 10% fetal bovine serum (FBS, heat inactivated) (Sigma), 0.1 mM MEM non-essential Amino Acids, 6 mM L-glutamine, 1mM MEM sodium pyruvate, and antibiotics (Gibco). For generation of VLP containing plasmid of interest (scramble, GFP or shRNA targeting SAMHD1), the day before transfection, cells were seeded on 10cm culture plates to reach 70% confluence for transfection. To produce transfection reagent, 9,6 µg of plasmid of interest was prepared together with 6µg Δ8.91 plasmid and 2,4 µg pCMV-VSV-g plasmid in DMEM complemented with 1/25 TransIT-293 transfection reagent (Mirus- Euromedex). The transfection reagent was then added drop by drop to HEK293FT. For generation of VLP containing Vpx (VLP/Vpx), HEK293FT were transfected by calcium- phosphate. The day before transfection, cells were seeded on 15cm culture plates to reach 70% confluence for transfection. Transfection reagent was composed of 50 µg pSIV3+ (Vpx coding plasmid^62^) with 10 µg pCMV-VSV-g plasmid in H20 + 0,25 M CaCl2 buffered by HBS according ProFection Mammalian Transfection protocol (Promega). Transfection reagent for VLP/Vpx was added drop by drop to HEK293FT. After one day, media of all transfected HEK293T cells (both for Vpx or plasmid of interest) was replaced with fresh medium. At day 2, supernatant was harvested and filtered twice using 0.45µm sterile filters (Sigma). VLP were immediately used for monocytes transduction.

#### Lentiviral transduction

Monocytes were transduced immediately after magnetic sorting. Medium was complemented with 8µg/ml polybrene (Sigma) to facilitate transduction, and cells were plated in 10cm culture plates. 5x10^6^ cells/mL of CD14^+^ in 5ml of complemented medium were transduced with 2,5 mL of VLP/Vpx and 4 mL of VLP containing plasmid coding shSAMHD1, scramble or GFP as described previously ^59^. Scramble transduced MDDC present unaffected transcriptomic profile and served as a point of reference for analysis. GFP coding plasmid allowed an assessment of transduction efficiency by flow cytometry. MDDC transduction efficiency > 90% at day 6 was considered satisfactory.

#### Flow cytometry analysis

For flow cytometry analysis, MDDCs were washed with PBS and first stained with LIVE/DEAD Fixable dye (TermoFisher) in PBS for 15 min at 4°C in the dark. After washing in PBS, cells were stained with antibodies for extracellular labelling for 25min at 4°C in PBS 1% Bovine Serum Albumin (BSA, Roche) and 1 mM EDTA (Life technology). For intracellular staining (ISG15), cells were fixed with cytofix/cytoperm (BD) for 15 min at room temperature then stained with antibodies in permeabilization buffer for 45 min at 4°C. After a wash, cells were resuspended in PBS and processed on BD fortessa flow cytometer. Analyses were performed of FlowJo software.

#### Seahorse XF Analyser experiment

The metabolic assays on MDDC cell culture were performed as previously described ^63^. Briefly, 100,000 cells were harvested at day 5 after transduction and plated in culture medium in 96- well Seahorse plates (Agilent). At day 6, cells were balanced for 1 h in unbuffered XF assay media (Agilent) supplemented for OCR analysis with either 2 mM Glutamine, 10 mM Glucose and 1 mM Sodium Pyruvate or just 2 mM Glutamine for ECAR measurement. For OCR measurements, compounds were injected during the assay at the following final concentrations: Oligomycin (ATP synthase inhibitor, 1 μM), FCCP (uncoupling agent measuring the maximal respiration capacity; 1 μM), Rotenone and Antimycin A (ETC inhibitors; 1 μM). For ECAR measurements, Glucose (10 mM), Oligomycin (1 μM), and 2-Deoxyglucose (2-DG, glycolytic inhibitor; 500 mM) were injected. For each cell line 4 to 6 technical replicates were evaluated. All OCR and ECAR measurements were normalized to the protein concentration dosed at the end of every experiment.

#### Quantitative RT–PCR

RNA was extracted from 100,000 cells following High Pure RNA isolation kit (Roche) instructions. RNA amplification was performed using Taqman primer probes and PCR master mix (TermoFisher) according to the manufacturer’s instructions or, for *CLPP* and *Hsp60* transcripts, the fast superior cDNA synthesis with SuperScript IV Vilo Reverse Transcriptase (Invitrogen) ^64^. Samples were analyzed on Viia7 Real-Time PCR system (ThermoFisher) and final data were plotted as expression relative to *HPRT*.

#### Western Blot

1 million cells were lysed in 100 µL of lysis buffer, composed of protease inhibitor, phosphatase inhibitor and NP40 (Sigma) in MiliQ Water, for 30 min on ice. Concentration of proteins were determined by Pierce BCA assay (ThermoFisher). After centrifugation clearance, 30 µg of protein lysates were boiled at 95°C for 5min in DTT and loading buffer (Invitrogen). Lysates were deposit on Bolt SDS page gel (Invitrogen) and transferred on nitrocellulose membrane (ThermoFisher) using Trans Blot Turbo transfert (Biorad). Membranes were saturated with TBS (Euromedex) 0.05% Tween20 (SIGMA) 5% BSA (Euromedex). Membranes were stained with indicated antibodies in TBS 0.05% Tween20 and ECL signal was recorded on Chemidoc (Biorad). Images were analyzed and band were quantified on ImageJ software.

#### Single-cell RNA sequencing

Frozen PBMC of AGS patients or healthy donors (CTRL) were processed by the LabTech single-cell@Imagine facility for cell encapsulation and NGS (Next generation Sequencing). A first experiment was performed on PBMC from healthy donors C1 and C2 and on PBMC from patients P2 before treatment and P1. A second experiment was performed on PBMC from healthy donors C3 and another vial from C1 (C1_bis) and on PBMC from another vial of P2 before treatment (P2_before_T_bis) and PBMC from P2 after treatment. The scRNA-seq libraries were generated using Chromium Single Cell 3′ Library & Gel Bead Kit v.2 (10x Genomics) according to the manufacturer’s protocol. Briefly, cells were counted, diluted at 1,000 cells/µL in PBS+0,04% and 20,000 cells were loaded in the 10x Chromium Controller to generate single-cell gel-beads in emulsion. After reverse transcription, gel-beads in emulsion were disrupted. Barcoded complementary DNA was isolated and amplified by PCR. Following fragmentation, end repair and A-tailing, sample indexes were added during index PCR. The purified libraries were sequenced on a Novaseq (Illumina) with 26 cycles of read 1, 8 cycles of i7 index and 98 cycles of read 2. Both experiments were integrated together on integration#1, thus including C1, C1_bis, C2, C3, P1, P2_before_T, P2_before_T_bis and P2_after_T.

Later, a new experiment was performed on PBMC from two healthy donors C4 and C5 as well as AGS patients with RNASEH2B mutation (P3 and P4), ADAR1 mutations (P5 and P6) and COPA patients (P7 and P8). This experiment account for integration#2. The scRNA-seq libraries were generated using Chromium Single Cell 3′ Library & Gel Bead Kit v.3 (10x Genomics) according to the manufacturer’s protocol. The purified libraries were sequenced on a Novaseq (Illumina) with 28 cycles of read 1, 8 cycles of i7 index and 91 cycles of read 2. During analysis, P1, P2_before_T, P2_before_T_bis and P2_after_T were often grouped together and labelled as “AGS5” group. Similarly, P3 and P4 were grouped as “AGS2”, P5 and P6 grouped as “AGS6” and P7 and P8 grouped as “COPA”. The integrations1# and #2 could not be merged due to technical advances (kit 10x for library generation version 2 versus kit version 3) made between experimental setup that do not allow direct comparison. Essentially, integration#2 allow significantly more genes to be detected. To assess RTI treatment effect on transcriptomic profile of P2 in **fig6 and S6**, P2_before_T_bis and P2_after_T were extracted from integration#1. These data were integrated together with data from healthy donors of the same experimental procedure C1_bis and C3 to generate integration#3 (see also supplementary table SI1).

#### Bioinformatic analysis

Sequenced reads from libraries generated were demultiplexed and aligned to the human reference genome (hg38), by Institut Imagine bioinformatic facility, using CellRanger Pipeline V6.0. R version 4.1.2 was used for data processing. Quality control, data integration and downstream analyses were produced using Seurat v4 ^65^. Apoptotic cells and empty sequencing droplets were removed by filtering cells with low features (nfeatures<250 for integration#1 and nfeatures<500 for integration#2) or a mitochondrial content <20%. Data were normalized using sctransform. For all three integrations generated, the Seurat clustering resolution 1.6 was selected for cluster identification. Labelling of cell types was defined based on expression of curated list of marker genes previously defined ^66^. Pathway enrichment were performed by applying the indicated list of DEG on EnrichR software ^23^. DEG were selected if fold change > 0.25 and false discovery rate ≤0.05. Enrichment of pathways in the list of DEG were ranked based on combined score. Combined score is a combination of adjusted *p*-value and z-score. *p*- value is computed using a standard statistical method used by most enrichment analysis tools: Fisher’s exact test or the hypergeometric test. This is a binomial proportion test that assumes a binomial distribution and independence for probability of any gene belonging to any set. z-score is computed using a modification to Fisher’s exact test in which we compute a z-score for deviation from an expected rank. EnrichR has lookup table of expected ranks and variances for each term in the library. These expected values are precomputed using Fisher’s exact test for many random input gene sets for each term in the gene set library. Enrichr uses this lookup table to calculate the mean rank and standard deviation from this expected rank as the z-score.

#### Gene regulatory network (GRN) inference analysis

To infer GRN, we apply pySCENIC (Single-Cell rEgulatory Network Inference and Clustering) ^16^ to a compendium of single-cell data from 8 AGS, 2 COPA patients and 6 healthy controls. Firstly, the gene modules that are co-expressed with transcription factors are identified using GRNboost^17^. Secondly, for each co-expressed module, the predicted transcription factor binding motifs and candidate transcription factors for a gene list are identified using *cis-regulatory* motif analyses using *RcisTarget*. To build the final regulons, we merge the predicted target genes of each TF-module that show enrichment of any motif of the given TF. Finally, regulator-target relationships are extracted and emitted as a set of network edges. Following the above pipeline, we obtain GRN for each sample. To further investigate the TF activity in AGS and control samples collectively, the GRNs of similar samples were integrated using similarity network fusion (SNF) ^67^ algorithm. The network-fusion step of SNF uses a non-linear method based on message passing theory ^68^ that iteratively updates every network, making it more similar to the others with every iteration. After a few iterations, SNF converges to a single network. The final network is analyzed and the top 50 TFs in AGS and control GRN were obtained by measuring out-degrees. To understand loss or gain of TF activity in AGS, we then performed differential out-degree analysis (DOA) ^69^. DOA scores for the top 50 TFs were obtained by taking the log2 ratio of the number of targets in AGS on the number of targets in control. TFs are then ranked based on absolute value of DOA score.

#### Cytokine measurement

Supernatant of the MDDC were obtained after centrifugation to remove debris at day 6 after transduction. Cytokines were measured using LEGENDplex Human Anti-Virus Response Panel (Biolegend) following the manufacturer’s protocol. Supernatant were not diluted before quantification. Fluorescence was measured on a BD fortessa cytometer.

### Quantification and statistical analysis

Statistical analyses were performed on Prism 8.0 (Graphpad) to calculate two-way ANOVA with Sidak’s multiple comparisons test or Wilcoxon rank test as indicated. ***:*p*<0.001, **:*p*<0.01, *:*p*<0.05. The number of unique donors for MDDC generation are listed in figure legends and represent biological replicates. Experiment with lower than 2 replicates were not statistically tested.

## Supporting information

SI1_Table1_Metadata_table

SI2_Table2_Signatures_gene_list

SI3_Seahorse_XF_analyser_time_course

SI4_DEG_lists

## Acknowledgements

This work was supported by the “*Institut national de la santé et de la recherche médicale*” (INSERM), the Atip-Avenir and “*Emergence ville de Paris*” programs and Fond de Dotation, Janssen Horizon and government grants managed by the *Agence Nationale de la Recherche* (ANR) as part of the “investment of the Future” program (Institut Hospitalo-Universitaire Imagine grant ANR-10-IAHU-01, Recherche Hospitalo-Universitaire grant ANR-18-RHUS-0010). The Labtech Single-Cell@Imagine is supported by the Paris Region and the “*Investissement d’avenir*” program through 2019 ATF funding – Sésame Filières PIA (grant 3877871). C.C. is the recipient of a CIFRE PhD (Sanofi). G.P.V. obtained an Imagine international PhD fellowship program supported by the Fondation Bettencourt Schueller. P.B. is the recipient of an ANRS post-doctoral fellowship. We thank the Imagine genomics, bioinformatics, cytometry and single-cell core facilities as well as metabolic analysis platform from Necker research institut. Patients’ cells were obtained according the research protocol CPP (ID-RCB/ EUDRACT: 2014-A01017-40) and we acknowledge referent physicians, Pr Marie Wisley (Cochin hospital, Paris), Dr Pierre Meyer (Neuropédiatrie, CHU Montpellier), Dr Florence Renaldo (Neuropédiatrie, CHU Trousseau, APHP), Dr Claude Cances Neuropédiatrie, CHU Toulouse), Pr Nadia Nathan (Pneumopédiatrie, CHU Trousseau, APHP), Pr Marie Wislez (Pneumologie, CHU Cochin, APHP), Dr Virgine Levrat (Pédiatrie, CH Annecy). We want to thanks all patients and their families.

## Author contribution

Conceptualization, M.B., M.M.M..; Methodology, M.B., M.M.M.; Formal Analysis, M.B, M.L., S.J., C.C.; Investigation, M.B., I.N., T.F., V.G.P., F.C.; Resources, B.N., M.L.F., P.Q.M., M.H., B.B.M, A.B., Y.C.; Writing - Original Draft, M.B., M.M.M.; Writing – Review and Editing, M.B., M.M.M., B.P., M.L.F., A.L., Y.C., A.F.; Visualisation, M.B., M.L., S.J., C.C.; Supervision, M.M.M., A.F.; Founding Acquisition, M.B, M.M.M.

## Supplemental information

Table SI1: Metadata table. Supplementary informations on human samples used.

Table SI2: List of genes constituting the different transcriptomic signatures used. Including references for each gene associated to mitochondrial stress in our signature and publicly available gene lists used in the analysis.

Fig SI3: Seahorse XF Analyzer time course including drug used and explanation on the measurement of glycolysis, max glycolytic capacity, ATP synthase activity and max respiratory capacity.

Table SI4: Differentially express gene (DEG) between AGS and CTRL groups listed by cell types.

## Data and code availability

• The scRNA-seq data discussed in this publication have been deposited in NCBI’s Gene Expression Omnibus (GEO) ^70^ and are accessible through GEO Series accession number GSE220764 (https://www.ncbi.nlm.nih.gov/geo/query/acc.cgi?acc=GSE220764).

• Any additional information required to reanalyze the data reported in this paper is available from the lead contact upon request.

---

**Fig S1:**
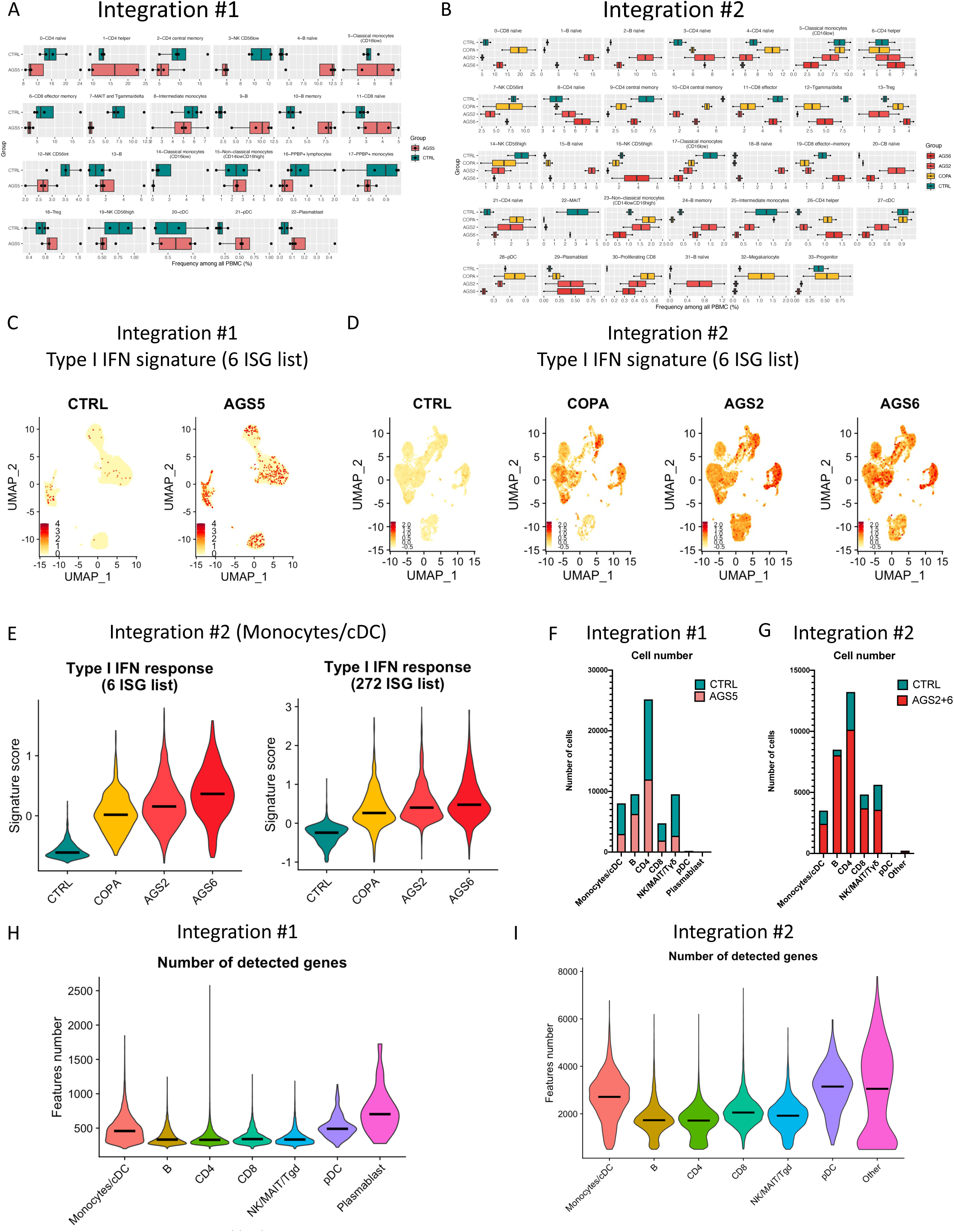
Quality control of cluster proportions, ISG signatures and DEG generated in all PBMC. Related to fig1. **(A and B)** Percentage of cell in indicated clusters among all PBMC between each group (CTRL, AGS2, AGS5, AGS6 and COPA) of integration#1 **(A)** and integration#2 **(B)** related to fig1E and 1F. All clusters are displayed. **(C and D)** Feature plot of ISG signature score of 6 ISG (*RSAD2, IFI27, IFI44L, ISG15, IFIT1, SIGLEC1*) (from Hadjadj at al. 2020) in integration#1 **(C)** and in integration#2 **(D)**. UMAP is split by CTRL, AGS2, AGS5, AGS6 and COPA groups and relate to fig1G and 1H. (E) Violin plot of type I IFN signature using the 6 ISG list (left panel) or the 272 ISG list (right panel) on monocytes/cDC of integration#2 split by CTRL, AGS2, AGS6 and COPA groups. Medians are shown as black lines. Related to fig1H. (F and G) Bar graph of the number cells in each cell types of integration#1 (F) and integration#2 (G) related to fig1I and 1J. (H and I) Violin plot of number of genes detected split by cell type from integration#1 (H) and from integration#2 (I). Medians are shown as black lines. In integration #1, cell types regroup several clusters at 1.6 resolution: “Monocytes/cDC” (clusters 5, 14, 17, 8, 20, 15), “B” (clusters: 4, 9, 10, 13,), “CD4” (clusters 0, 1, 2, 16, 18), “CD8” (clusters 6, 11), “NK/MAIT/Tγδ” (clusters 3, 7, 12, 19), pDC (cluster 21) and “Plasmablast” (cluster 22). In integration#2 cell types regroup several clusters at 1.6 resolution: “Monocytes/cDC” (clusters 5, 17, 23, 25, 27), “B” (clusters: 1, 2, 15, 18, 20, 24, 31), “CD4” (clusters 3, 4, 6, 8, 9, 10, 13, 21, 26), “CD8” (clusters 0, 11, 19, 30), “NK/MAIT/Tγδ” (clusters 7, 12, 14, 16, 22), pDC (cluster 28) and “Others” (cluster 29, 32, 33). cDC= classical Dendritic cells; B= B lymphocytes; CD8= Lymphocytes T CD8^+^; CD4= Lymphocytes T CD4^+^; NK= Natural Killer cells; MAIT= Mucosal-associated invariant T cells; Tγδ= Lymphocytes Tγδ; pDC= plasmacytoid dendritic cells.

**Fig S2:**
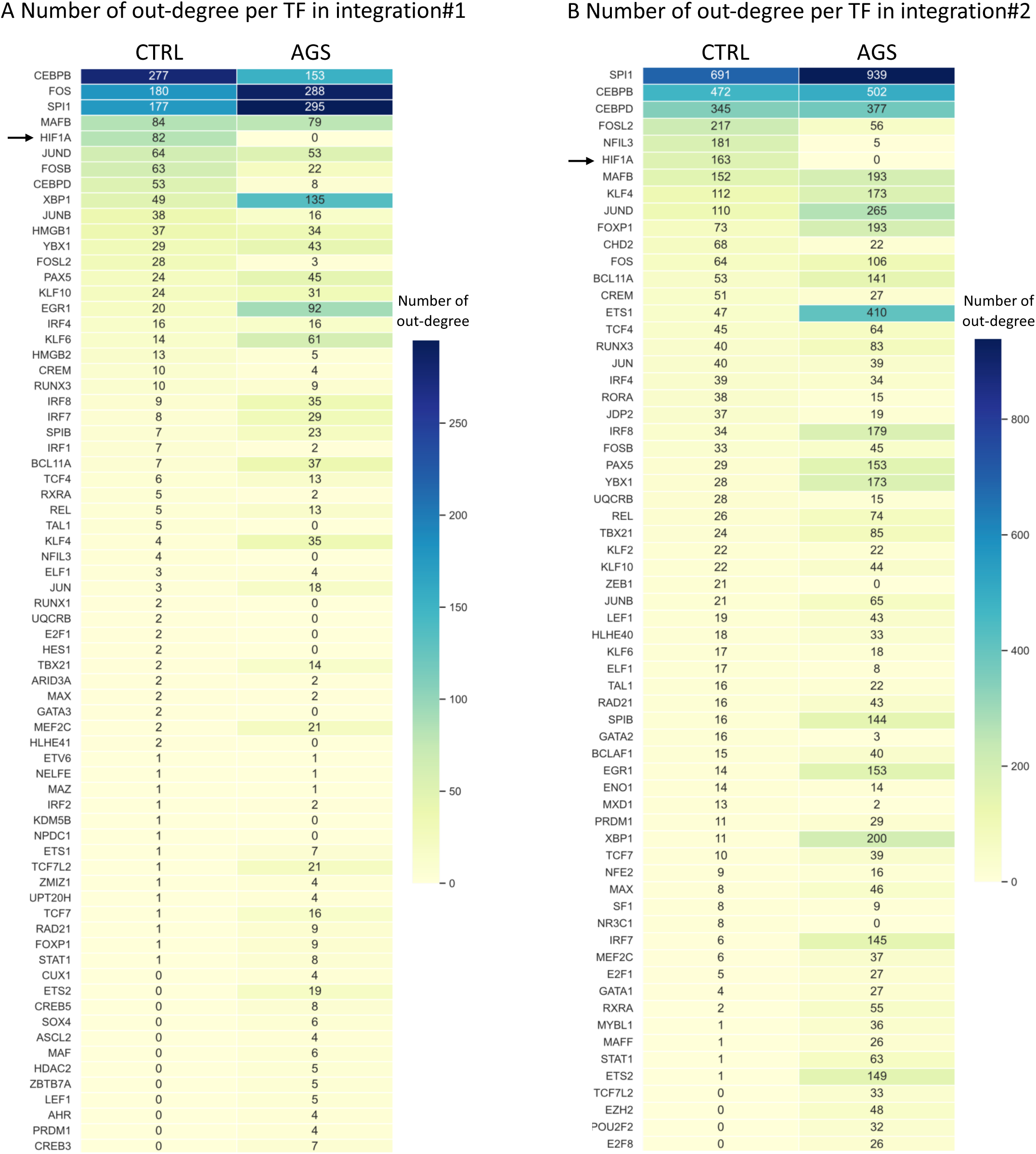
HIF-1α activity is predicted to be important in shaping transcriptomic profile of healthy PBMC in CTRL. Related to fig2. (A and B) Heatmap of out-degree score of top50 TF in CTRL and corresponding score in AGS patients of integration#1 (A) and integration#2 (B) used for differential out-degree analysis in fig2B and 2D. Black arrow points HIF-1α.

**Fig S3:**
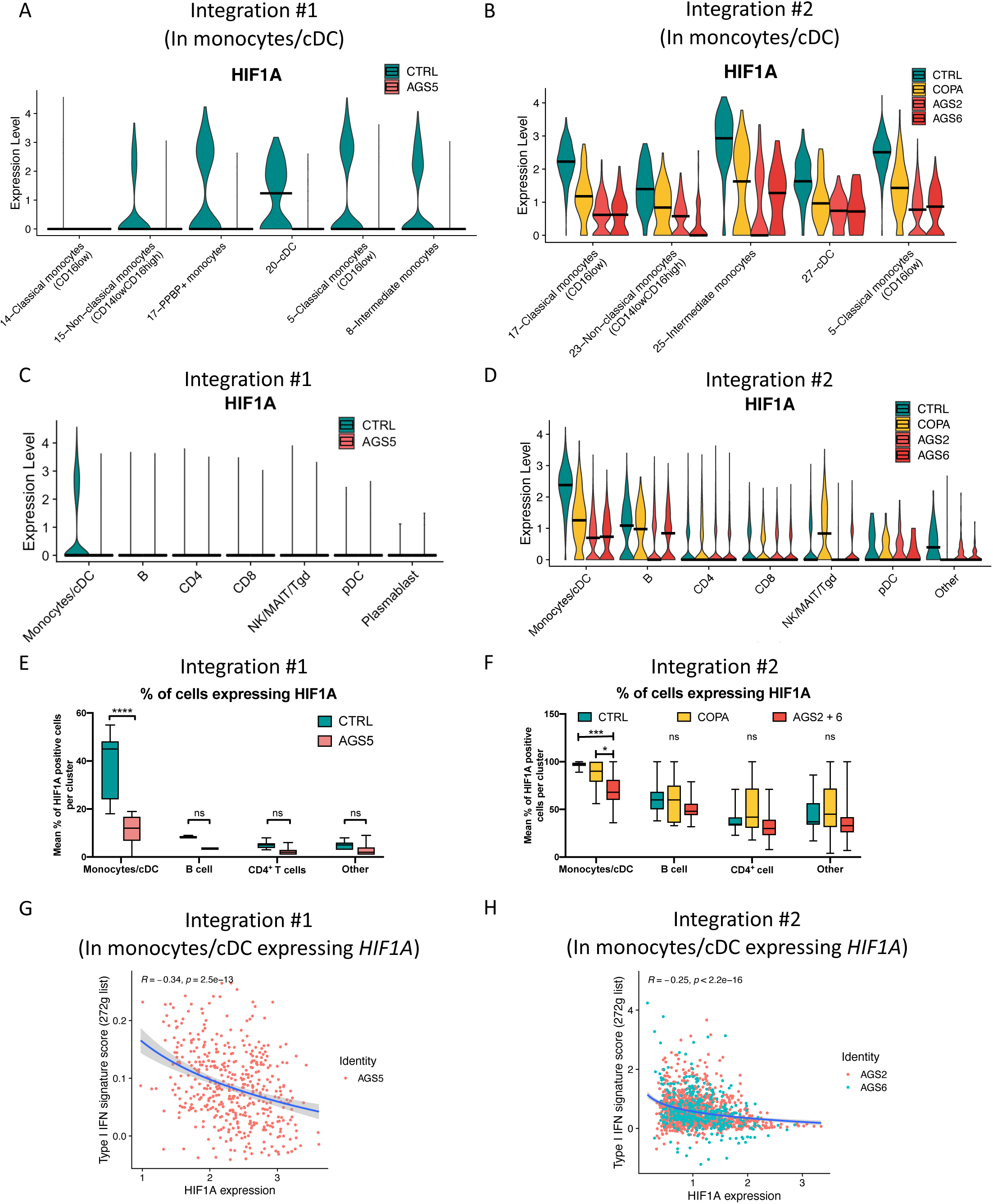
*HIF1A* loss of expression in AGS is stronger in monocytes/cDC than in any other cell type. Related to fig3. (A and B) Violin plot of *HIF1A* expression in each cluster belonging to monocytes/cDC split by groups from integration#1 (A) and from integration#2 (B) related to fig3C and 3D. Medians are shown as black lines. (C and D) Violin plot of *HIF1A* expression in each cell type split by CTRL, AGS2, AGS5, AGS6 and COPA groups from integration#1 (C) and from integration#2 (D) related to fig3C and 3D. Medians are shown as black lines. **(E and F)** Percentage of cells expressing *HIF1A* mRNA in each cluster grouped by cell type in CTRL, AGS2, AGS5 and AGS6 groups from integration#1 **(E)** and integration#2 **(F)**. Medians, 10-90% quartiles and min to max are represented as box plot. One way anova with Sidak multiple correction statistical test are represented as: ***:*p*<0,001, **:*p*<0,01, *:*p*<0,05, ns: not significant. (G and H) Correlation of *HIF1A* expression with type I IFN signature score (272 genes list) in monocytes/cDC of patients from integration#1 (G) and integration#2 (H). Only *HIF1A* expressing cells are displayed, each point is a cell and colors discriminate mutated form of AGS. In integration #1, cell types regroup several clusters at 1.6 resolution: “Monocytes/cDC” (clusters 5, 14, 17, 8, 20, 15), “B” (clusters: 4, 9, 10, 13,), “CD4” (clusters 0, 1, 2, 16, 18), “CD8” (clusters 6, 11), “NK/MAIT/Tγδ” (clusters 3, 7, 12, 19), pDC (cluster 21) and “Plasmablast” (cluster 22). In integration#2 cell types regroup several clusters at 1.6 resolution: “Monocytes/cDC” (clusters 5, 17, 23, 25, 27), “B” (clusters: 1, 2, 15, 18, 20, 24, 31), “CD4” (clusters 3, 4, 6, 8, 9, 10, 13, 21, 26), “CD8” (clusters 0, 11, 19, 30), “NK/MAIT/Tγδ” (clusters 7, 12, 14, 16, 22), pDC (cluster 28) and “Others” (cluster 29, 32, 33). cDC= classical dendritic cells; B= B lymphocytes; CD8= Lymphocytes T CD8^+^; CD4= Lymphocytes T CD4^+^; NK= Natural Killer cells; MAIT= Mucosal-associated invariant T cells; Tγδ= Lymphocytes Tγδ; pDC= plasmacytoid dendritic cells.

**Fig S4:**
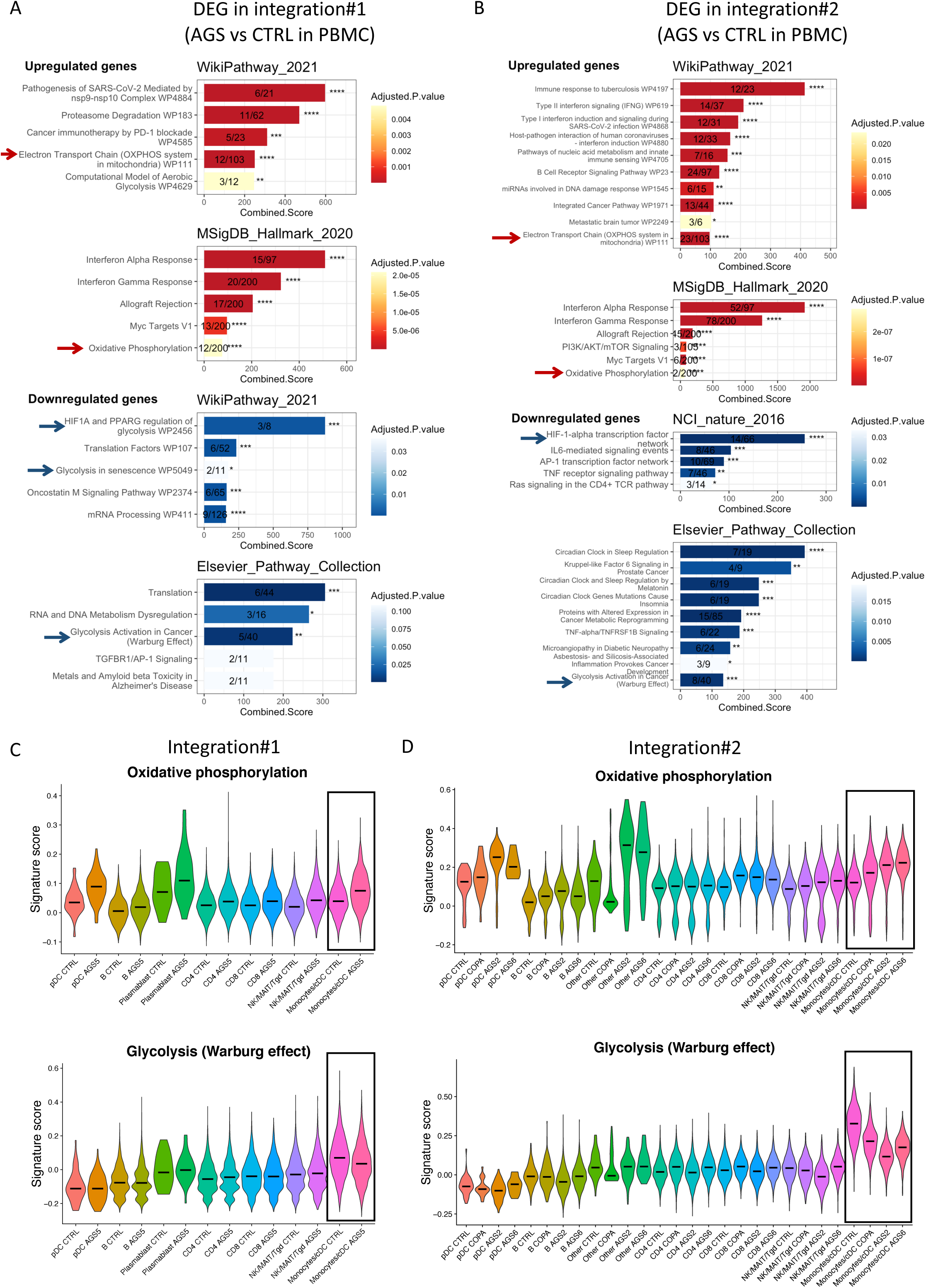
Energy metabolism switch is also observed while analyzing all PBMC but is more noticeable in monocytes/cDC. Related to fig4. **(A and B)** Pathway enrichment analysis using EnrichR (Chen et al. 2013) of DEG in AGS patients (AGS5) compared to CTRL of integration#1 **(A)** or in AGS patients (AGS2 + AGS6) compared to CTRL of integration#2 **(B)**. Pathways are ranked based on combined score. Color gradient is proportional to adjusted *p-*value of Fisher’s test and labelled as ***:*p*<0.001, **:*p*<0.01, *:*p*<0.05. Numbers indicates the number of genes present in the DEG list over the total number of genes referenced in the corresponding pathway. Red highlight pathways including genes upregulated in patients and blue highlight pathways including genes downregulated. Red arrows point toward oxidative phosphorylation associated pathways and blue arrows point toward anaerobic glycolysis and HIF-1α-related pathways. (C and D) Violin plot representing signature score of “Oxidative phosphorylation pathway” (MSIgDB hallmark datatbase) (Top panel) and of “Glycolysis Activation in Cancer Warburg Effect” (Elsevier database) (Bottom panel) on PBMC split by cell types as well as CTRL, AGS2, AGS5, AGS6 and COPA groups from integration#1 (C) and integration#2 (D). Medians are shown as black lines. Squares indicates monocytes/cDC. Refer to fig4. In integration #1, cell types regroup several clusters at 1.6 resolution: “Monocytes/cDC” (clusters 5, 14, 17, 8, 20, 15), “B” (clusters: 4, 9, 10, 13,), “CD4” (clusters 0, 1, 2, 16, 18), “CD8” (clusters 6, 11), “NK/MAIT/Tγδ” (clusters 3, 7, 12, 19), pDC (cluster 21) and “Plasmablast” (cluster 22). In integration#2 cell types regroup several clusters at 1.6 resolution: “Monocytes/cDC” (clusters 5, 17, 23, 25, 27), “B” (clusters: 1, 2, 15, 18, 20, 24, 31), “CD4” (clusters 3, 4, 6, 8, 9, 10, 13, 21, 26), “CD8” (clusters 0, 11, 19, 30), “NK/MAIT/Tγδ” (clusters 7, 12, 14, 16, 22), pDC (cluster 28) and “Others” (cluster 29, 32, 33). cDC= classical dendritic cells; B= B lymphocytes; CD8= Lymphocytes T CD8^+^; CD4= Lymphocytes T CD4^+^; NK= Natural Killer cells; MAIT= Mucosal-associated invariant T cells; Tγδ= Lymphocytes Tγδ; pDC= plasmacytoid dendritic cells.

**Fig S5:**
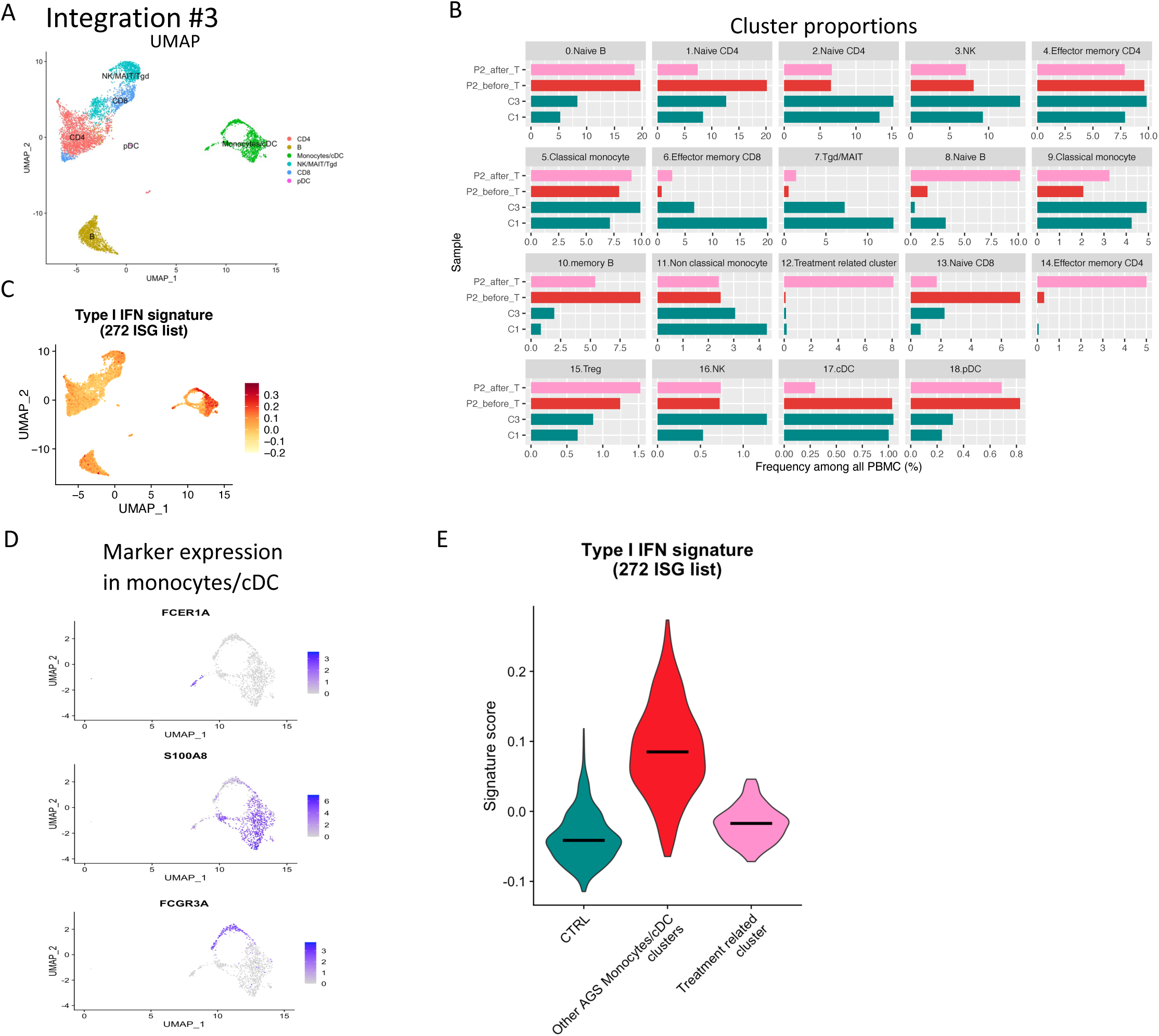
Characterization of the treatment-related cluster. Related to fig5. (A) UMAP of integration#3 including label of main cell types related to fig5A. (B) Cell proportion of each cluster from the UMAP resolution 1.6 among PBMC of integration#3 related to fig5B. (C) Feature plot of type I IFN signature score (272 ISG list) on PBMC of integration#3 related to fig5B. (D) Feature plots of *FCER1A, S100A8* and *FCGR3A* expressions in monocytes/cDC of integration#3 used for respective identification of cDC, classical monocytes and non-classical monocytes related to fig5C. (E) Violin plot of ISG signature score (272 genes list) on monocytes/cDC from integration#3, related to fig5G. Median are shown as black lines. Related to fig5B. In integration#3, “Monocytes/cDC” refers to clusters 5, 9, 11, 12, 17 at 1.6 resolution. In figS5, clusters are grouped as follows: “Treatment related cluster” (cluster 12 from P2_after_T), “Other AGS Monocytes/cDC clusters” (clusters 5, 9, 11, 17 from both P2_before_T and P2_after_T samples). cDC= classical dendritic cells; B= B lymphocytes; CD8= Lymphocytes T CD8^+^; CD4= Lymphocytes T CD4^+^; NK= Natural Killer cells; MAIT= Mucosal-associated invariant T cells; Tγδ= Lymphocytes Tγδ; pDC= plasmacytoid dendritic cells.

**Fig S6:**
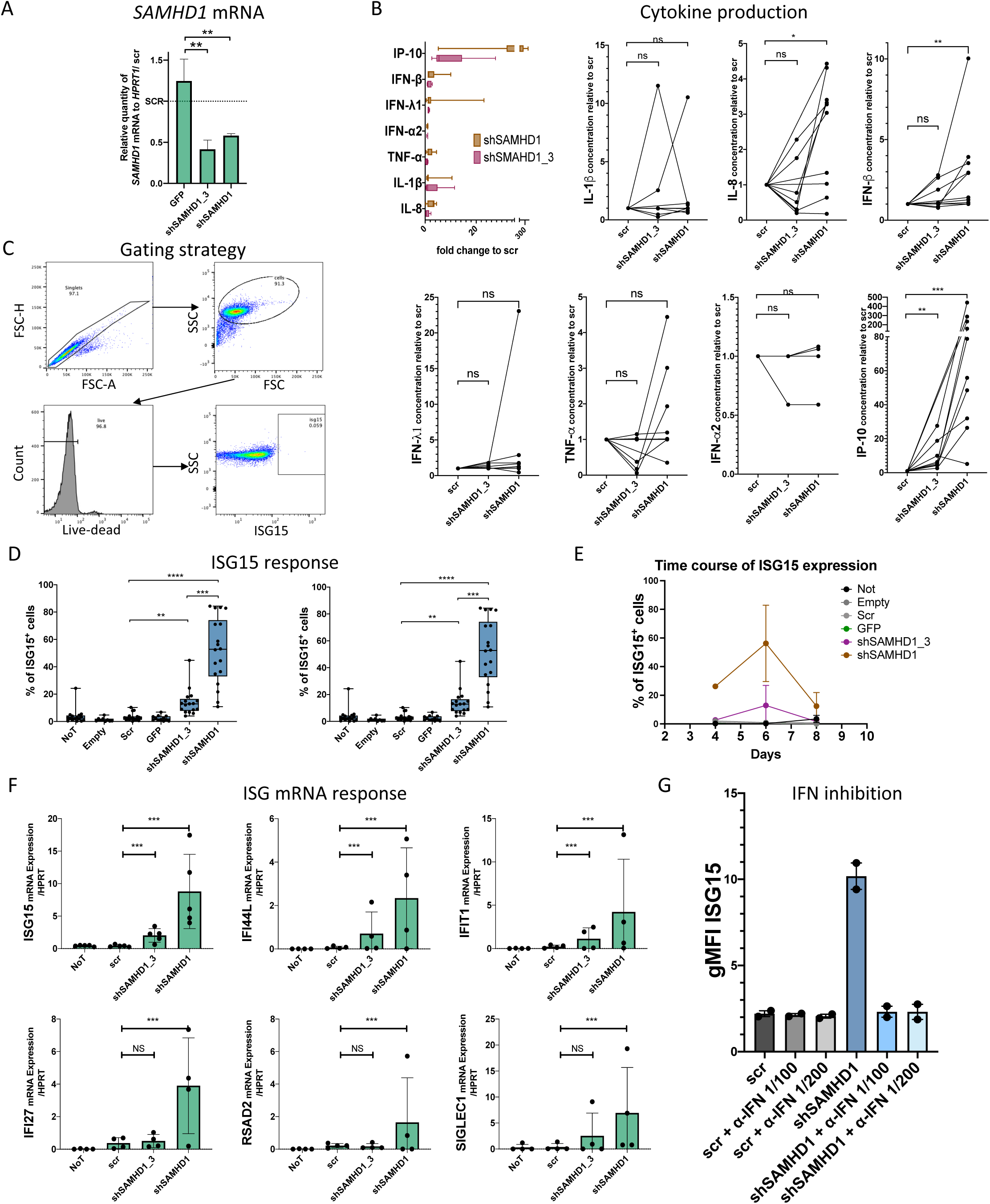
Results observed in our AGS *in vitro* cellular model are reproduced using a second shRNA targeting SAMHD1. Related to fig6. MDDC were transduced by another shRNA targeting *SAMHD1* mRNA (shSAMHD1_3) and results are compared to the shRNA presented in fig6 (shSAMHD1) (A) Refers to fig6A but with the addition of shSAMHD1_3. Anova’s statistical test is represented as: ns: not significant, ***:*p*<0.001, **:*p*<0.01, *:*p*<0.05. (B) Refers to fig6C but with the addition of shSAMHD1_3. All cytokines tested are presented as fold change of supernatant concentration to scramble condition, all cytokines together (top left panel) or individually (other panels). Wilcoxon’s statistical test against scr group is represented as: ns: not significant, ***:*p*<0.001, **:*p*<0.01, *:*p*<0.05. (C) Flow cytometry gating strategy referring to fig6D. Representative plot of isotype control for ISG15 staining is shown. (D) Refers to fig6E with the addition of shSAMHD1_3 (Left panel). Geometric mean fluorescent intensity (gMFI) of ISG15 staining is also represented (right panel). (E) Time course of ISG15 expression tracked by flow cytometry on transduced MDDC from day 4 to day 8 after transduction. Means with standard deviations form triplicate for 1 healthy donor are represented. (F) mRNA level of each individual genes used for score calculation in fig6F. mRNA level of each gene was quantified by RT-qPCR and normalized to *HPRT1* mRNA level on MDDC generated from 5 unrelated healthy donors. Refers to fig6F with the addition of shSAMHD1_3. (G) gMFI of ISG15 intracellular staining by flow cytometry on shSAMHD1 transduced MDDC in presence of type I IFN blocking antibodies from 2 unrelated healthy donors. Refers to fig6K. α-IFN=Blocking antibody cocktail against type I IFN. **(D and F)** Means with SEM and one way anova with Sidak multiple correction statistical test are represented as: ns: not significant, ***:*p*<0.001, **:*p*<0.01, *:*p*<0.05. (A to G) NoT: Not transduced MDDC; Empty: MDDC transduced with an empty vector; scr: MDDC transduced with a scramble RNA without known target in human cells; GFP: MDDC transduced with a GFP coding plasmid for assessment of transduction efficiency; shSAMHD1_3: MDDC transduced with the shRNA against *SAMHD1* n°3; shSAMHD1: most efficient shRNA targeting *SAMHD1* represented in main fig6.

**Fig S7:**
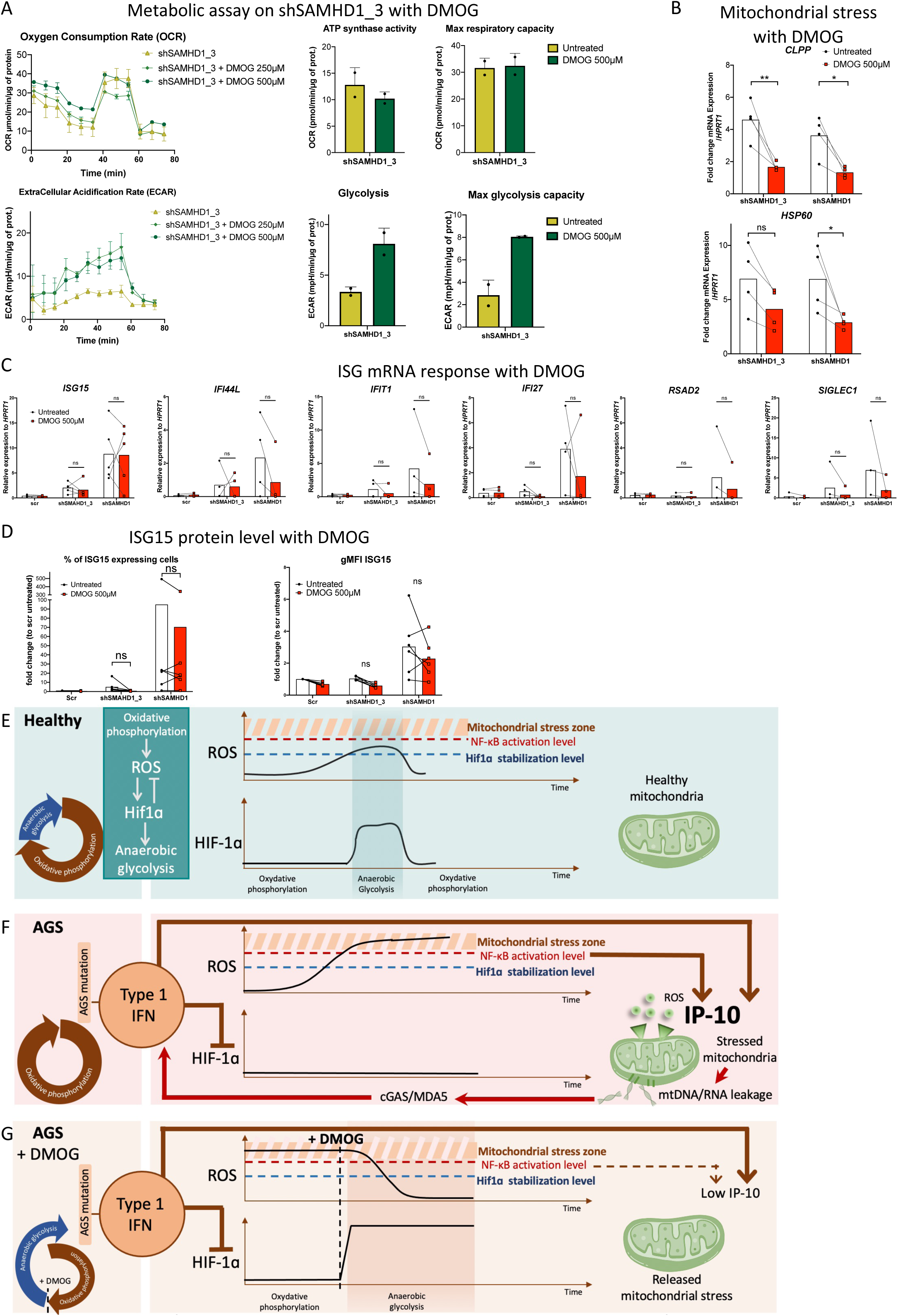
Proposed molecular model in AGS pathogenesis. Related to fig7. MDDC were transduced by another shRNA targeting *SAMHD1* (shSAMHD1_3) and results are compared to the shRNA presented in fig7 (shSAMHD1). (A) MDDC at day 4 after transduction were treated with increasing concentration of DMOG for 48h before undergoing Seahorse XF Analyzer time course for oxygen consumption rate (OCR) (top panels) or ECAR (bottom panels). A Seahorse XF Analyzer time course is represented for 1 healthy donor for both assays. Mean with standard error of triplicates are represented (left panels). ATP synthase activity, Max respiratory capacity, actual glycolysis and max glycolytic capacity were quantified for 4 unrelated healthy donors assessed by Seahorse XF Analyzer experiment (right panels). Related to fig7B. **(B)** RT-qPCR of *CLPP* and *HSP60* on RNA extracted from MDDC at day 5 after transduction (24hours of DMOG) for 4 unrelated healthy donors. Data were normalized to *HPRT1* gene. Related to fig7C. (C) mRNA quantification of individual genes used for score calculation in fig7D. mRNA level of each gene is normalized to *HPRT1* mRNA level from 5 unrelated healthy donors. (D) Percentage of ISG15 expressing cells (left panel) and gFMI of ISG15 on MDDC at day 6 after transduction (48hours of DMOG) by flow cytometry on 6 unrelated healthy donors. Refer to fig7E. (A to E) shSAMHD1_3: MDDC transduced with the shRNA against SAMHD1 n°3; shSAMHD1: most efficient shRNA targeting *SAMHD1* represented in main fig7. **(B, C and D)** Mean and one way anova with Sidak multiple correction statistical test are represented as: ns: not significant, *:p<0,001, **:p<0,01, ***:p<0,05 (E, F and G) We propose a model combining our observations and current literature detailed in discussion. (E) HIF-1ɑ is known to senses then controls ROS production through oxidative phosphorylation pathway by balancing energy metabolism toward anaerobic glycolysis. **(F)** In AGS patients, we observed loss of HIF-1ɑ activity and expression associated with an energy metabolism locked in oxidative phosphorylation state, aggravated mitochondrial stress and excessive IP-10 production. We showed that HIF-1ɑ stabilisation prevent part of IP-10 production independently of type I IFN. IP-10 transcription is known to be induced following type I IFN signaling but can also be induce by NF-κB signaling. Accordingly, we propose that, without HIF-1ɑ, uncontrolled ROS production by excessive oxidative phosphorylation might be an NF-κB activator and participate to elevated IP-10 production in patients. Mitochondrial stress might be a source of ligands for the nucleic acid sensing pathway fuelling type I IFN production. **(G)** We showed that *in vitro* treatment using DMOG on an MDDC model of AGS, restored HIF-1ɑ activity, reverted energy metabolism, released mitochondrial stress and reduced IP-10 production. ROS= Reactive oxygen species; HIF-1ɑ= Hypoxia inducible factor 1ɑ; NF-κB= Nuclear factor κB; AGS= Aicardi Goutières syndrome; IP-10= Interferon inducible protein 10; mtDNA/RNA= mitochondrial (deoxy)ribonucleic acids; DMOG= dimethyloxalylglycine.

