## Supplementary material for "Enhanced inflammatory signaling driven by metabolic switch in Aicardi-Goutières syndrome": SI3_Seahorse_XF_analyser_time_course

Protocol 1: Mitochondria activity profile (Phosphorylation oxidative)

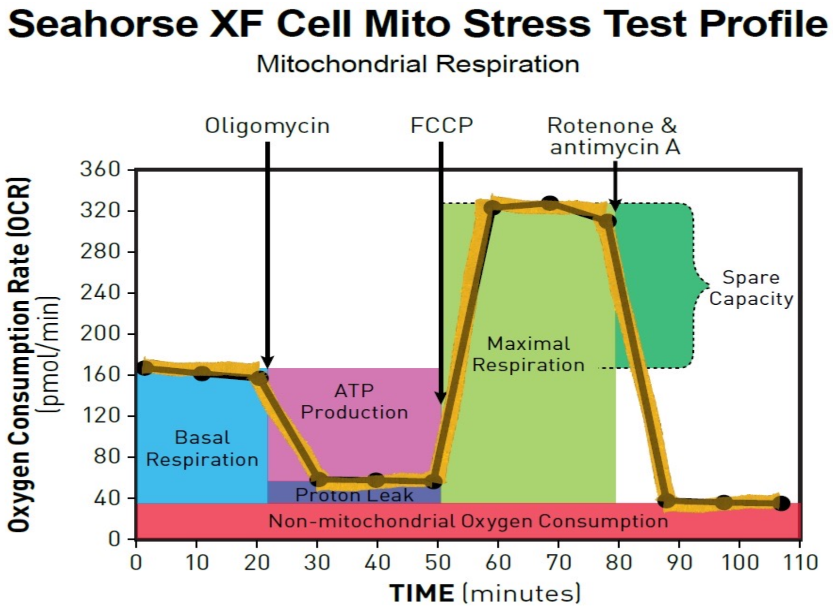


Drug addition at indicated time points

**Oligomycin**: supress ATP synthase

**FCCP**: Uncouple ATP synthase activity from oxygen consumption

**Rotenone & antimycin**: supress all the electron transport chain

Protocol 2: Glycolysis profile

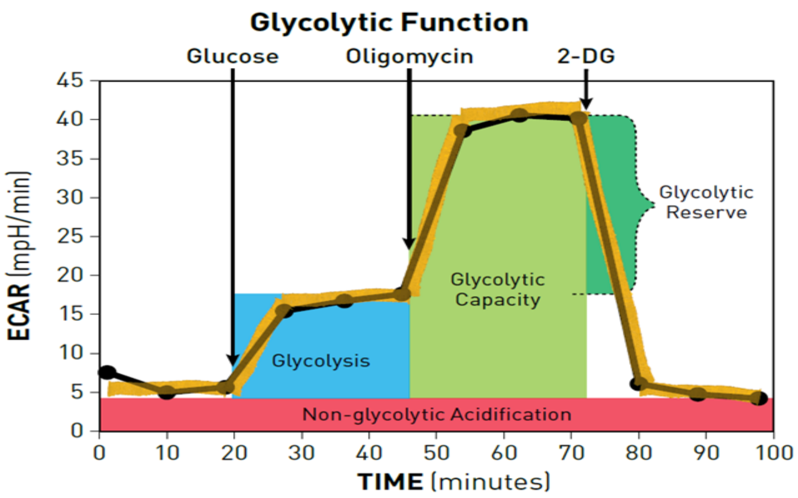


ECAR: extracellular consumption rate

**Glucose**: allows glycolysis

**Oligomycin**: supress ATP synthase

**2-DG**: supress the first step of glycolysis
